# CDC25B couples ribosome biogenesis to cell cycle kinetics to promote spinal cord neurogenesis

**DOI:** 10.64898/2026.10.29.762835

**Authors:** Sofiane Omerani, Laurence Liaubet, Simon Lebaron, Celia Plisson-Chastang, Fabienne Pituello, Sophie Bel-Vialar

## Abstract

During neurogenesis, neural progenitors undergo a tightly regulated transition from proliferation to differentiation, which requires precise coordination between cell cycle progression and molecular reprogramming. Although the role of CDC25B is well established in promoting G2/M transition through CDK1 activation, we previously showed that it is both necessary and sufficient to trigger neural progenitor differentiation, via CDK-dependent and CDK-independent mechanisms. However, the molecular networks underlying this neurogenic activity remain poorly understood. Here, using a gain-of-function approach in the chicken spinal cord combined with bulk RNA sequencing, we show that CDC25B overexpression upregulates transcripts encoding ribosomal proteins and enhances ribosome biogenesis (RiBi) in neural progenitors. Consistent with these findings, during normal development, RiBi increases in neurogenic progenitor domains expressing CDC25B. Notably, the increase in RiBi is not associated with an increase in translation in neural progenitors. Instead, it is followed by an increase in mTORC1 activity in neuronal precursors and subsequently by a rise in the global translation rate in differentiating neurons. Moreover, increasing mTORC1 activity is sufficient to trigger a premature transition of neuronal precursors to the next step of differentiation. Altogether, our data support a model in which CDC25B coordinates cell cycle progression with RiBi in neurogenic progenitors, thereby initiating a program that culminates in translation in differentiating neurons.

## INTRODUCTION

Coordinating progenitor expansion with timely differentiation is a fundamental requirement for building and maintaining tissues. In the developing central nervous system (CNS), deregulation of this balance underlies a wide range of human neurodevelopmental disorders, from microcephaly or spina bifida, to various psychiatric conditions. In the developing spinal cord, different sets of neurons and glial cells are sequentially produced from proliferating progenitors located into the neural tube. Differentiation begins in the ventral region of the neural tube with the production of motor neurons and a subset of interneurons, then it gradually extends to the dorsal region where dorsal sensory interneurons are produced (Frith et al., 2024; Sagner and Briscoe, 2019). Shortly after their birth close to the lumen of the neural tube, neuronal precursors migrate laterally out of the ventricular zone and accumulate in the mantle zone, where they differentiate into post-mitotic neurons. These different steps are controlled by proneural genes such as NEUROG2 or MASH1, which coordinates cell cycle exit with differentiation, reinforced by positive feedback mechanisms that precede the onset of pan-neuronal genes (Bertrand et al., 2002; Lacomme et al., 2012; Gui et al., 2007; Novitch et al., 2001; Diez del Corral and Storey, 2001). How individual progenitors integrate these signals and decide, division after division, whether to self-renew or to commit to neuronal differentiation remains a central question.

At the cellular level, this choice involves progressive changes in the mode of division of neural progenitors (Götz and Huttner, 2005). Progenitors first perform self-expansive proliferative divisions that generate two progenitors (PP), while being specified according to their dorso-ventral position through the action of morphogens such as BMP/Wnt dorsally and Sonic hedgehog (Shh) ventrally (Morin et al., 2007; Sagner and Briscoe, 2019; Saade et al., 2013). They then progressively switch to a neurogenic mode of division, in which at least one daughter cell differentiates into a neuron. It occurs through either asymmetric self-replacing divisions generating a progenitor and a neuron (PN), or symmetric terminal divisions, producing two neurons (NN) (Morin et al., 2007; Wilcock et al., 2007). This transition is accompanied by changes in cell cycle kinetics: Although proliferation rate is spatially uniform across progenitors, the length of individual cell cycle phases correlates with neurogenesis (Dalton, 2015; Molina and Pituello, 2017). In proliferative domains, progenitors display short G1 and long G2 phase duration, whereas G1 is longer and G2 shorter in neurogenic domains (Peco et al., 2012; Kuzmicz-Kowalska and Kicheva, 2020). In addition, as the neural tube matures, the total cell cycle length increases with time, mainly through G1 lengthening (Molina and Pituello, 2017), thereby reducing the proliferative capacity of neural progenitors (Azaïs et al., 2019; Kuzmicz-Kowalska and Kicheva, 2020). Accordingly, core regulators of the cell cycle machinery coordinate cell cycle progression with neurogenesis. For instance, in the neural tube CyclinD1 regulates cell proliferation but also promotes neurogenesis, independently of its interaction with the cell cycle machinery (Lobjois et al., 2008; Lukaszewicz and Anderson, 2011). Cdkn1c, a cyclin-dependent kinase inhibitor, has also recently emerged as a key regulator of the transition from symmetric proliferative to asymmetric neurogenic divisions (Mida et al., 2025). Cell cycle regulators can also modulate proneural activity, as demonstrated with NEUROG2, whose activity is controlled through quantitative phosphorylation of serine/proline sites by CDK1/2 (Ali et al., 2011; Hindley et al., 2012; Hindley and Philpott, 2012). We previously showed that the G2/M regulator CDC25B, a phosphatase expressed in neural progenitors, also displays a neurogenic activity, consistent with the shorter G2 phase observed in neurogenic domains (Boutros et al., 2006; Peco et al., 2012; Bonnet et al., 2018; Roussat et al., 2023). CDC25B expression correlates temporally and spatially with neuronal differentiation, and, at the onset of neuronal differentiation, is restricted to the ventral neural tube (Peco et al., 2012). CDC25B downregulation in the ventral domain lengthens G2 phase and impairs neuronal production, whereas CDC25B gain-of-function in dorsal progenitors shifts the balance from proliferative toward neurogenic divisions, both self-replacing and terminal, resulting in increased neuronal differentiation (Bonnet et al., 2018). Notably, a modified form, CDC25B^ΔCDK^, which cannot interact with CDK1, still promotes self-replacing neurogenic divisions but fails to enhance terminal ones, demonstrating that part of CDC25B pro-neurogenic activity is independent of its canonical cell cycle function (Bonnet et al., 2018). Collectively, these findings identify the cell cycle machinery as a central node in the regulation of neuronal fate acquisition dependent or independent of their cell cycle activity. However, how cell cycle regulators influence fate decisions beyond their canonical function remains largely unknown.

In addition to their action through transcriptional regulation, another level at which such regulation could operate is the control of the protein abundance of fate determinants. Indeed, emerging data highlight the important role of translational regulation in refining the level of molecular determinants that trigger neuronal determination and differentiation (Zahr et al., 2019). Cell cycle regulators themselves are under such control; although Cdkn1C transcripts are uniformly expressed, low protein levels initially regulate the progressive transition of progenitors into neurogenic modes of division before higher expression later drives cell cycle exit in daughter cells (Mida et al., 2025). In the chick spinal cord, heterogeneous PAX6 protein levels, despite uniform transcript expression, determine neuronal commitment and differentiation (Bel-Vialar et al., 2007). In the developing cortex, NEUROG2/TUBB3 transcripts are present in apical progenitors while their proteins appear only in intermediate progenitors and neurons (Telley et al., 2016). In adult tissues, Sox2 translation is progressively repressed as neural stem cells transition toward a neurogenic and neuroblast state (Baser et al., 2019). Together these examples underscore the critical role of temporal control of protein synthesis downstream of transcriptional regulation as a means of rapidly adjusting protein abundance in response to internal and external signals (Li and Wang, 2020).

The global translational capacity of a cell is ultimately set by its ribosome content. Ribosomes, composed of rRNA and ribosomal proteins, are assembled in the nucleolus through multiple coordinated biological processes collectively referred to as ribosome biogenesis (RiBi) (Baßler and Hurt, 2019; Henras et al., 2015). The importance of ribosome homeostasis for neural development is highlighted by ribosomopathies, a group of disorders caused by RiBi defects whose CNS manifestations include microcephaly, autism and neural tube closure defects. (Aubert et al., 2018; Bourque et al., 2018; Herrlinger et al., 2019; Hetman and Slomnicki, 2019; Farooq et al., 2020; Wang et al., 2023). Likewise, deregulation of the mTOR pathway, a positive regulator of RiBi and translation, causes neurodevelopmental and neurodegenerative disorders (Lipton and Sahin, 2014), as do mutations in translational machinery components such as elongation and initiation factors (Yang et al., 2014; Zahr et al., 2018; Kolaj et al., 2023). Moreover, RiBi is dynamically regulated during corticogenesis (Chau et al., 2018). These findings establish ribosome homeostasis as a critical determinant of neuronal development and support translation as a key trigger of the proliferative-to-neurogenic transition. Importantly, RiBi is tightly coupled to cell cycle progression in other cellular contexts: the nucleolus disassembles at mitotic entry and reassembles at mitotic exit under the control of CDK activity (Sirri et al., 2002; Hernandez-Verdun, 2011), and rRNA synthesis is repressed in mitosis by CDK1/cyclin B-mediated phosphorylation of the Pol I transcription machinery, while being activated in G1 by G1-specific CDK–Cyclin complexes (Heix et al., 1998; Voit et al., 1999; Voit et al., 2015; Iyer-Bierhoff and Grummt, 2019). Thus, both cell cycle kinetics and ribosome homeostasis are closely linked to neurogenesis. Yet how ribosome synthesis is coordinated with cell cycle kinetics to control cell fate commitment during neurogenesis remains poorly understood.

Here, using a combination of high throughput sequencing, functional analyses and electron microscopy in the developing chick spinal cord, we identified a link between the cell cycle machinery and the regulation of RiBi during spinal neurogenesis. We found that CDC25B enhances RiBi in neurogenic progenitors and this is consistent with our observation of a highly dynamic regulation of RiBi and translation rate accompanying neuronal commitment and differentiation. We also provided evidences that increasing the mTORC1 pathway is sufficient to accelerate neuronal differentiation. Together, these results support a model in which CDC25B couples cell cycle progression to neuronal differentiation by controlling both cell cycle kinetics and RiBi.

## RESULTS

### Differential analysis of the CDC25B transcriptome reveals significant enrichment of pathways related to ribosome biogenesis and translation

To identify the main molecular events associated with progenitors’ proliferation to differentiation switch driven by CDC25B, we combined a gain of function (GOF) strategy using *in ovo* electroporation, with a global measure of transcripts levels using bulk RNA sequencing. To be able to distinguish between pathways specifically modified through the interaction of CDC25B with CDK1 from those regulated independently, we compared GOF of CDC25B (hCDC25B) with the mutated form of CDC25B that prevents CDC25B–CDK1 interaction (hCDC25B^ΔCDK^). This CDC25B variant does not display any effect on G2 phase length while conserving CDC25B phosphatase activity (Bonnet, Molina et al. 2018). We overexpressed hCDC25B, hCDC25B^ΔCDK^ or LacZ as a control, under the control of a CDC25B specific promoter (see method), together with H2B-GFP. Electroporated neural tubes were collected 24h post electroporation and dissociated cells were sorted to isolate GFP+ cells for RNA sequencing and analysis (**Fig. 1A**).

**Figure1:**
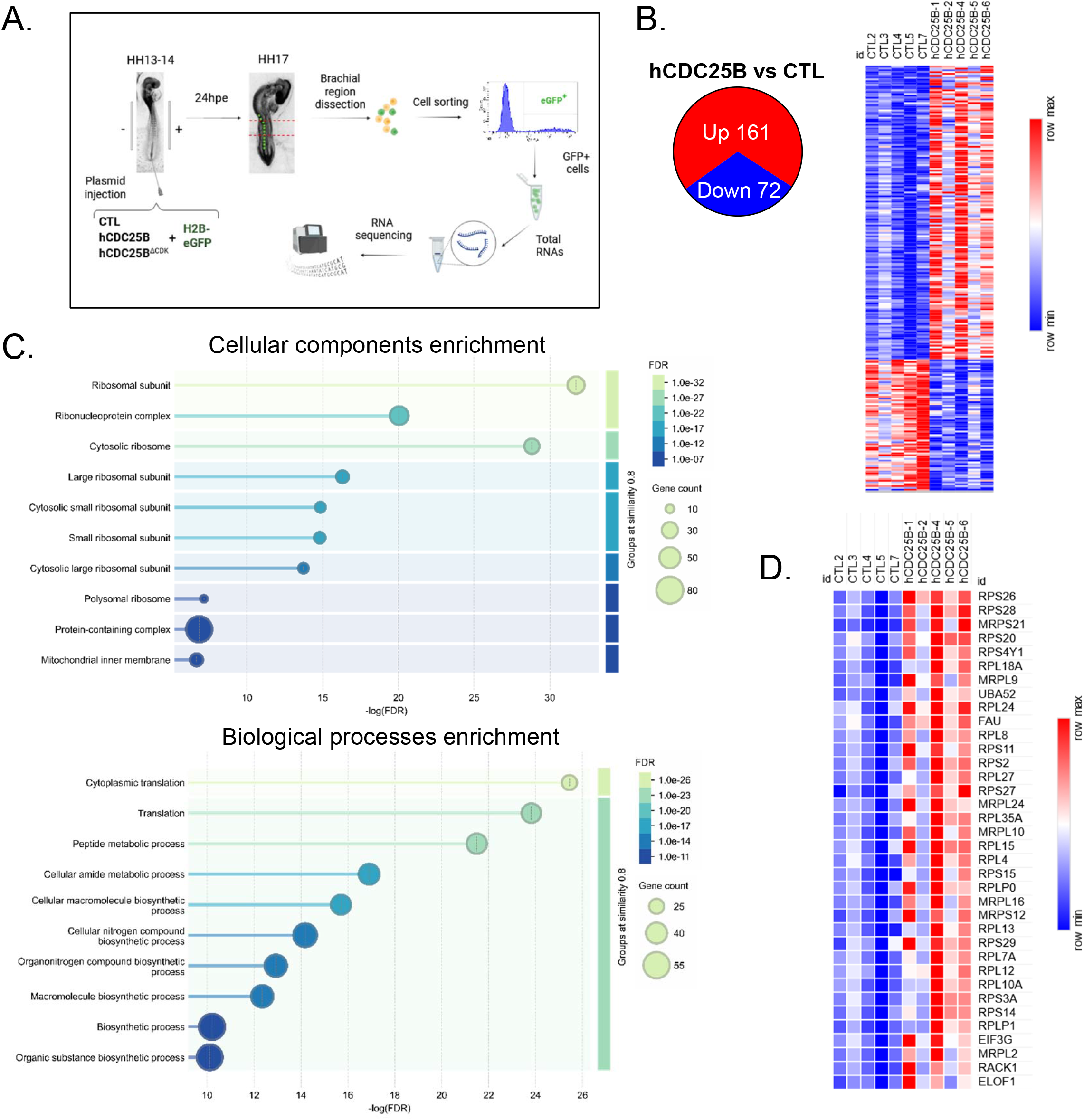
Differential analysis of the hCDC25B transcriptome reveals significant enrichment of pathways related to ribosome biogenesis and translation. (A) Schematic representation of RNA-Seq strategy. Experiments were performed on chicken embryos at stages HH13–14, co-electroporated with a control plasmid (pMP-LacZ) or a plasmid enabling overexpression of wild-type hCDC25B (pMP-hCDC25B) or hCDC25B^ΔCDK^ (pMP-hCDC25B^ΔCDK^) together with a plasmid encoding H2B-eGFP. Embryos were collected 24 hours post-electroporation (at stage HH17-18). Neural tubes were dissected from the brachial region (somites 12–20), and dissociated cells were sorted to enrich for GFP-positive cells expressing the respective constructs. Each experimental condition was independently replicated 6 times, with each technical replicate consisting of neural tubes pooled from 10 to 13 embryos, ensuring high reproducibility and robustness of the transcriptomic analysis. Total RNA was extracted with Tri reagent (Thermofisher) and sent to the Beijing Genomics Institute (BGI) for sequencing. 5 replicates were kept for each condition after selection based on RNA quantity and quality. (B) Left: Proportion of differentially expressed genes upregulated (red) and downregulated (blue) in hCDC25B condition relative to the control among the 233 genes identified with FDR < 0.1. Right: Heatmap illustrating the expression profiles of the 161 upregulated genes and the 72 downregulated genes in the hCDC25B condition compared to the control. (C) Gene Ontology (GO) enrichment analysis. Top 10 significantly enriched GO terms in the *cellular component* and *biological process* categories among the 161 upregulated genes, as determined by string-db. Enriched terms are ranked by FDR-adjusted p-value, with the x-axis indicating the - log(FDR) value. (D) Heatmap illustrating the genes associated with the GO terms “translation” with their expression levels in hCDC25B gain-of-function relative to the control condition. Genes are classified based on their fold change statistical significance (FDR).

The principal component analysis (PCA) showed that the hCDC25B and CTL samples are well segregated along the PCA1 axis whereas the hCDC25B^ΔCDK^-GOF samples (DelCDK) were not well grouped **(Fig.S1)**. We therefore started by performing differential analysis between CTL and hCDC25B conditions, without considering the hCDC25B^ΔCDK^-GOF samples. We identified 233 differentially expressed transcripts between the CTL and hCDC25B-GOF (FDR< 0.1) (**Fig. 1B**). Gene ontology (GO) term enrichment analysis reveals that the most enriched terms are related to ribosome and translation (**Fig. 1C**). Among the genes that are enriched following hCDC25B-GOF, we detected 24 genes coding for ribosomal proteins, 7 genes coding for mitochondrial ribosomal proteins and genes coding either for nucleolar protein and/or related to the translational machinery (**Fig. 1D**; FAU, UBA52, ELOF1, NIP7, IMP3, LSM10, EIF3G and ZNF593). This first analysis suggests that the main modification occurring downstream of CDC25B in neural progenitors is an enhancement of ribosome biogenesis.

### WGCNA analysis of bulk RNA-Seq data revealed a high correlation between ribosome, neurogenesis pathways and endogenous CDC25B expression

We further explored our raw data by conducting a *Weighted correlation network analysis* (WGCNA) to identify cluster of genes (modules) sharing correlated expression pattern. The objective was both to determine if ribosomal proteins were co-regulated and associated with specific gene networks/functions, and to explore how the different modules were correlated with the expression of CDC25B. As the gene clustering occurs independently of the experimental conditions, we performed this second analysis on all our samples (CTL, hCDC25B and hCDC25B^ΔCDK^) to increase robustness. A subset of 5000 genes with the highest variance across the 15 samples was selected. The WGCNA analysis allowed the identification of 13 modules corresponding to co-expression networks, containing 81 to 1675 genes (**Fig.2A**). Ribosomal proteins were grouped in a unique module, the blue one, indicating that they are co-regulated in the developing spinal cord.

**Figure 2:**
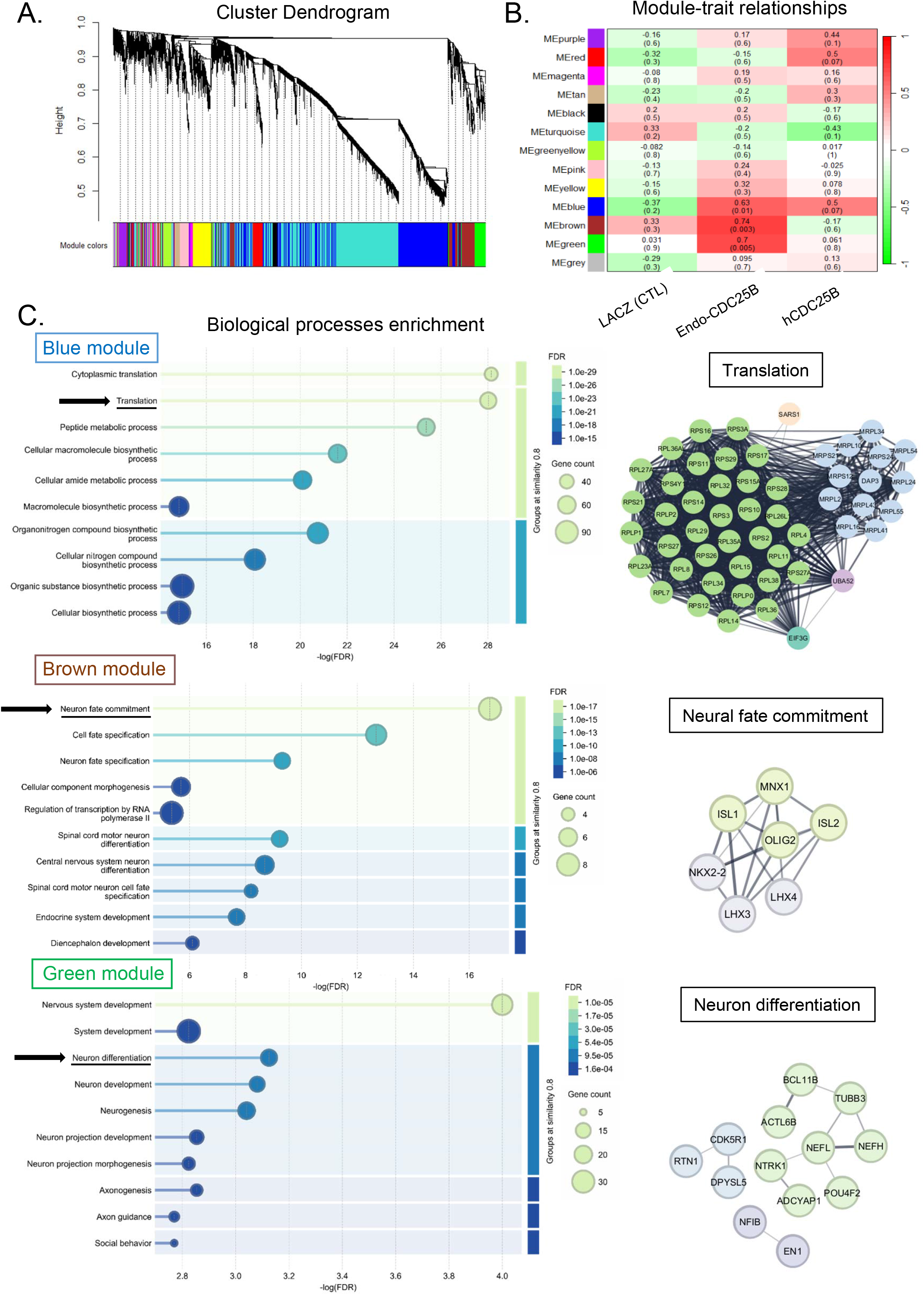
WGCNA analysis of bulk RNA-Seq data identified a high correlation of ribosome and neurogenesis pathways with endogen-CDC25B expression. (A) Clustering dendrogram of 5000 genes with the highest variance, with dissimilarity based on topological overlap, together with assigned module colors. (B) Trait-module associations between expressed genes and traits related to quantification of CDC25B. Each row corresponds to a module eigengene and each column to a trait. The corresponding correlation and p-value are shown in each cell. The table is color-coded according to the color legend, which shows the correlations. (C) Ontological analysis of the blue, brown and green modules selecting positive correlation and p-value<0.05 with endogen-CDC25B. On the left, graphs showing the representative top 10 biological processes enrichment for each module. On the right, highlight of relevant genes networks from biological process enriched in each module (arrows). “Translation” gene networks: in green genes coding for ribosomal proteins; in blue, genes coding for mitochondrial ribosomal proteins. “Neural fate commitment” gene network: in green genes related to motoneuron specification; in blue, genes related to ventral interneurons specification. “Neuron differentiation” gene network: In green and blue, two subnetworks related to neuronal differentiation.

In a second step, to identify co-regulation between gene modules and CDC25B expression, we quantified endogenous-CDC25B (chicken CDC25B), hCDC25B, and LacZ (expressed in the control vector) in each sample **(Fig.S2)** and used these expression levels as traits. The correlation analysis between each module and traits identified significant correlations (**Fig.2B**, p-value < 0.05) between endogenous-CDC25B and the blue module (1265 genes), the brown module (406 genes), and the green module (236 genes), of which, the blue module was also correlated with the hCDC25B gain of function (Corr>0.5, pval<0.07). No correlation was detected between the 13 modules and hCDC25B^ΔCDK^ or LacZ. We analysed the content of these three modules selecting positive correlation and p-value<0.05 with endogenous-CDC25B. Ontology analysis of the blue module reveals a significant correlation between ribosomal proteins and endogenous CDC25B (**Fig.2C**). Ribosomal protein networks are also significantly correlated with the hCDC25B-GOF **(Fig.S3)** confirming the result of our differential analysis. The brown and green modules contain gene networks related to ventral neuronal differentiation (e.g., *Islet1/2*, *Olig2*, *NKX2.2*, *LHX3/4*) and neuronal function (e.g., genes involved in nervous system development, neuronal differentiation, and axon guidance), respectively. This is coherent with the endogenous CDC25B expression in ventral progenitors at the stage we collected the neural tubes (**Fig.2C**).

This WGCNA analysis corroborates the data presented above showing that the hCDC25B-GOF increases ribosomal proteins transcription. Importantly, it also highlights a strong correlation between the endogenous CDC25B expression and genes related to RiBi /translation. Altogether, these correlations between CDC25B and enrichment of ribosomal proteins suggest their common association with neuronal differentiation.

### CDC25B gain-of-function increases ribosomal biogenesis

Our high throughput sequencing analyses indicate that ribosomal protein transcripts are upregulated downstream of CDC25B. In order to see if this upregulation is linked to an increase of ribosome biogenesis, we measured the nucleoli volumes, as a proxi for RiBi (Baker, 2013; Sanchez et al., 2016), 24h following electroporation of hCDC25B or a control vector together with GFP. We performed immunostaining for nucleophosmin (NPM1), a histone chaperone involved in the assembly of the 40S and 60S ribosomal subunits, to label nucleoli (**Fig. 3A**) and, measured nucleoli volumes in GFP and non GFP progenitors on Z-stack images using the Imaris software (Fig. 3B and see Methods). We detected significantly larger nucleoli in hCDC25B electroporated cells compared to CTL, whereas there was no difference between non-electroporated cells in the two conditions (**Fig. 3C**). Hence, CDC25B induces an increase in ribosome biogenesis, which is consistent with the increase in transcripts encoding ribosomal proteins downstream of CDC25B observed in our RNA-Seq experiments.

**Figure 3:**
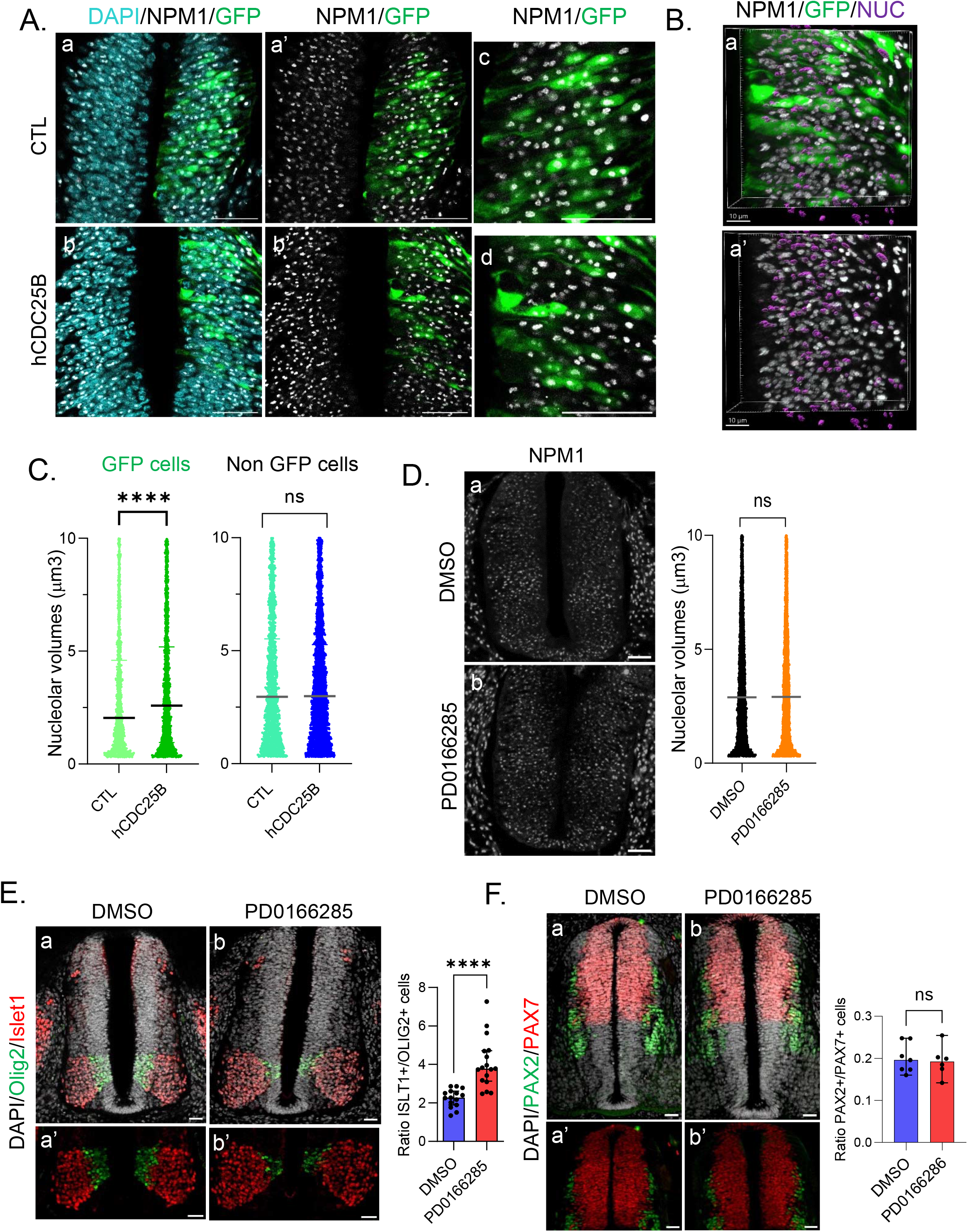
The increase of ribosomal protein transcription downstream of CDC25B is associated with increase ribosomal activity. (A) Confocal images of HH18 chick spinal cord, 24 hr after co-electroporating either pmp-lacZ (CTL, a, a’), or pmp-hCDC25B (b,b’) with pMIW-GFP and stained with NPM1. (c) Close up images of a, a’; (d) close up image of b,b’. Scale bars 30 µm. (B) 3D visualisation of a region from b obtained with Imaris software showing 3D reconstruction of nucleoli in purple. Scale bar 10µm. (C) Quantification of nucleolar volume (µm³) based on NPM1 immunostaining and 3D reconstruction in the dorsal neural tube of CTL or hCDC25B electroporated embryos in GFP cells or in the contralateral non-electroporated side (non-electroporated cells). Quantifications were performed across at least 3 sections from 3 embryos of three different experiments, with data expressed as median ± IQ range (each point representing the volume of a single nucleolus). Mann-Whitney test **** p< 0,0001, ns. Non-significant. Scale bars:30 µm. (D) Left: Representative cross-sections of HH18 chick spinal cord, 24 hr after DMSO or PD0166286 treatment, stained with NPM1 (white). Right: Nucleolar Volumes (µm3) quantifications were performed across at least 3 embryos, with data expressed as median (each point representing the volume of a single nucleolus). Mann-Whitney test, ns. Scale bars 30 µm. (E) Left: Transverse sections of the HH18 chicken spinal cord 48 hours following DMSO (a,a’) or PD0166286 (b,b’) treatment, immunostained with OLIG2 (progenitors) and ISLET1 (neurons). Right: Quantification of the percentage of progenitors’ vs neurons in the ventral part of the neural tube in the two experimental conditions with data expressed as median ± IQ range. Kruskal-Wallis test ****p< 0,0001. ns. Non-significant. Scale bars 30 µm. (F) Left: Transverse sections of the HH18 chicken spinal cord 48 hours following DMSO (a,a’) or PD0166286 (b,b’) treatment, immunostained with PAX7 (progenitors) and PAX2 (neurons). Right: Quantification of the percentage of progenitors’ vs neurons in the dorsal part of the neural tube in the two experimental conditions with data expressed as median ± IQ range. Kruskal-Wallis test ****p< 0,0001. ns. Non-significant. Scale bars 30 µm.

As CDK1 is the canonical CDC25B target for cell cycle regulation, we then tested if increasing CDK1 activity in neural progenitors was sufficient to trigger a RiBi increase. We treated HH12-13 embryos with PD0166285, a Wee1/Myt1 inhibitor (Baffet et al., 2015) for 24h *in ovo*. As Wee1 and Myt1 prevent mitosis entry through negatively regulating CDK1 activity (Potapova et al., 2009), their inhibition leads to an increase in CDK1 activity and to a shortening of the G2 phase duration (Roussat et al., 2023). We validated the efficiency and the specificity of our treatment by measuring cell cycles parameters of neuronal progenitors in treated vs untreated embryos. The G2 phase length is reduced upon PD0166285 treatment without significant effect on G1, S or cell cycle length **(Fig.S4)**. We then checked for RiBi by measuring nucleolar volumes and found that increasing CDK1 activity is not sufficient to trigger an increase of RiBi in progenitors (**Fig. 3D**). An explanation for this difference between CDC25B GOF and PD0166285 treatment could be that this pharmacological treatment is not sufficient to trigger neuronal differentiation. To test that hypothesis, we treated HH12-13 embryos with PD0166285 or DMSO and checked for neuronal differentiation 48h following treatment, in the dorsal and ventral part of the neural tube. In the ventral domain, we observed an increase in the number of Islet1+ neurons at the expense of Olig2+ progenitors in PD0166285 treated embryos but to lesser extend that what we had observed with the hCDC25B gain-of-function **(Fig3E)** (Bonnet et al., 2018). However, this treatment had no significant effect in the dorsal domain, as the ratio between PAX2+ dorsal neurons and PAX7+ dorsal progenitors is the same between samples treated with PD0166285 or DMSO **(Fig3F).**

These results indicate that increasing CDK1 activity may enhance neuronal differentiation in the CDC25B-expressing ventral domain, but is not sufficient to trigger RiBi and neuronal differentiation on its own in the non-neurogenic dorsal domain. This suggests that the CDC25B-induced increase in RiBi is part of its pro-neurogenic activity and may be, to some extent, independent of the cell cycle.

### Ribosome biogenesis is dynamically regulated during neurogenesis

The finding that CDC25B triggers an increase of RiBi and that CDC25B expression correlates with increased expression of ribosomal proteins prompted us to investigate how RiBi is modulated during the process of spinal neurogenesis. We analysed markers of ribosome biogenesis at two key developmental stages in the chick spinal cord: at HH12-13 (E2), when proliferating progenitors predominantly undergo proliferative divisions, and neuronal production begins marginally in the ventral domain (Le Dreau et al., 2014), and at HH18-19 (E3.5), when the cervico-brachial region of the neural tube is divided into two main domains: a dorsal region where progenitors mainly undergo proliferative divisions, and a ventral domain enriched in neurogenic progenitors that actively generate neurons. To assess RiBi activity, we immunostained neural tube sections with NPM1, which localizes to the granular component (GC), and fibrillarin (FBL), which is enriched in the dense fibrillar component (DFC), where it mediates early pre-rRNA processing (Zhang et al., 2024) and (**Fig. 4A**).

**Figure 4:**
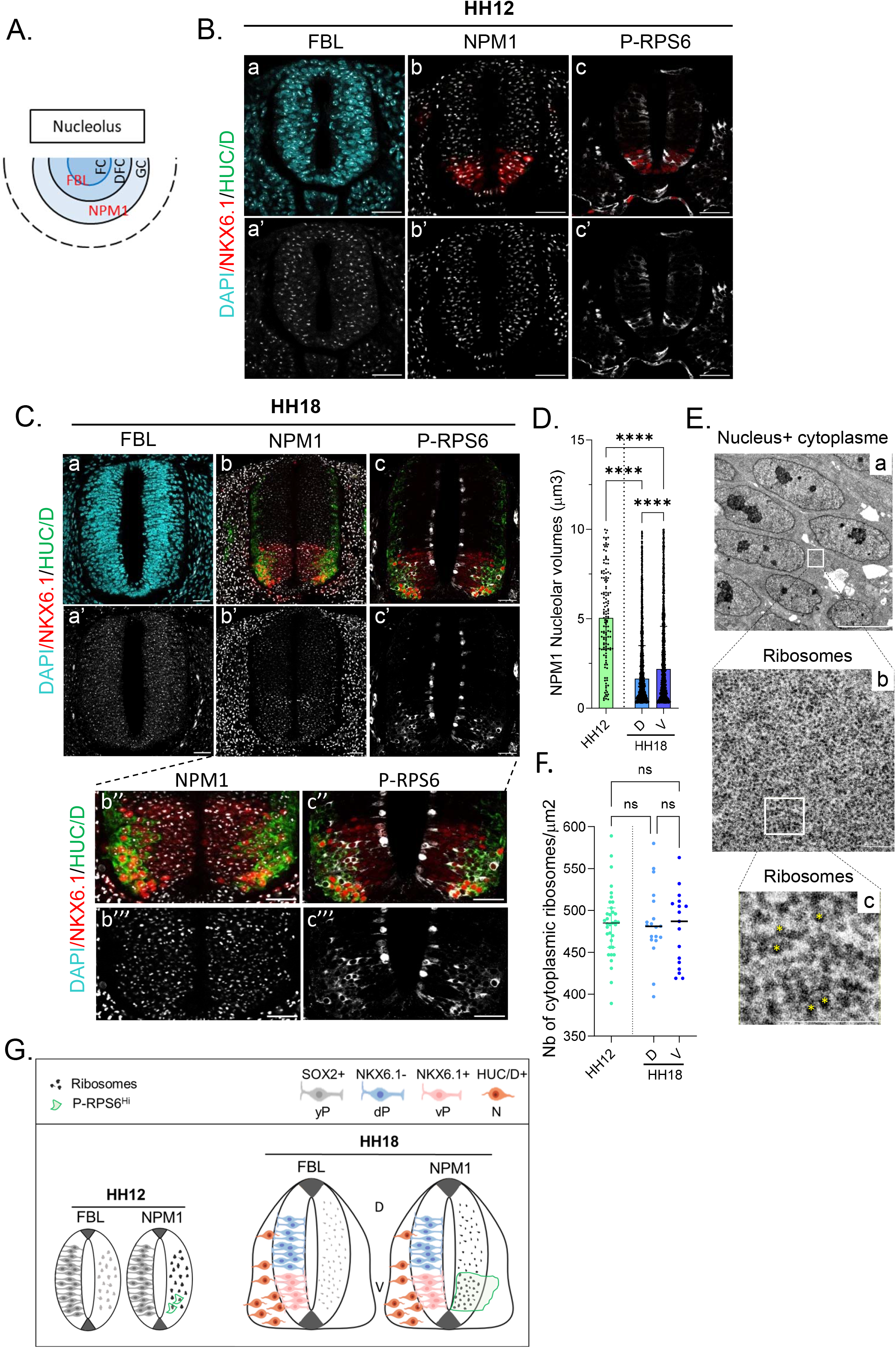
Dynamic of RIBI and translation during spinal neurogenesis. (A) Schematic representation of the nucleolus showing its three major domains: the fibrillar center (FC), dense fibrillar component (DFC), and granular component (GC). Fibrillarin (FBL) and nucleophosmin (NPM1) are shown as markers of the DFC and GC, respectively, highlighting the spatial organization of ribosome biogenesis within the nucleolus. (B) Transverse sections of the HH12 chicken spinal cord stained with DAPI (blue), and immunostained with NKX6.1 (red) and HuC/D (green, no staining at this stage) plus FBL (a–a′), NPM1 (b–b′) or phosphorylated RPS6 (P-RPS6, c–c′). Scale bars 30 μm. (C) Representative transverse sections of HH18 chicken spinal cord stained for DAPI (blue) and immunolabeled for NKX6.1 (red), HuC/D (green), together with FBL (a–a′), NPM1 (b–b′), P-RPS6 (c–c′). Bottom: Enlarged views of the ventral neural tube domain highlighting the distribution of nucleolar markers. Scale bars 30 μm. (D) Nucleolar volume (µm³) quantified from NPM1 immunostaining in the neural tube of HH12 embryos and in dorsal versus ventral domains at HH18. Quantifications were performed across at least 3 embryos, with data expressed as median ± IQ range (each point representing the mean of n ≥ 3 sections per embryo). Kruskal-Wallis test, **** p< 0,0001. Scale bars 30 µm. (E) Transmission electron microscopy (TEM) images of (a) neural progenitors in the ventricular zone at HH12 - nucleus containing nucleolus are visible -, (b) close up on ribosomes in perinuclear cytoplasmic regions and (c) enhanced view of ribosomes. Colour stars in the enlarged panels identify some individual ribosomes. Scale bars 200 nm. (F) Quantitative analysis of cytoplasmic ribosomes abundance in HH12 progenitors and in dorsal versus ventral progenitors at HH18. Quantifications were performed on images such as E(b) using ImajeJ across at least 3 embryos at each stage (see methods), with data expressed as median ± IQ range (each point representing the value of the counting in one cell). Kruskal-Wallis test, ns, non-significant. (G) Scheme summarising the temporal dynamic of RiBi in the neural tube based on our observations. At HH12 nucleoli are big and uniformly sized along the D-V axis. At HH18-19, nucleoli are smaller than at HH12 but there are larger in the ventral domain of the neural tube, a neurogenic region, than in the dorsal domain that does not yet actively produce neurons. Ventral neurogenic region also displays higher P-RPS6 staining (in green). yP (young neural progenitor), dP (dorsal progenitor), vP (ventral progenitor), and N (neuron).

At HH12-13, we observed homogenous expression of FBL and NPM1 in all the nucleoli along the dorso-ventral axis of the neural tube (**Fig. 4B**). At HH18-19, whereas the FBL staining still shows homogeneous pattern, the intensity of the NPM1 staining is now strongly diminished in the dorsal region of the neural tube while staying high in ventral progenitors, identified with NKX6.1 labelling (**Fig. 4C**). This progenitor domain expresses high levels of CDC25B and actively produces motoneurons at this stage (Peco et al., 2012). We then measured nucleoli volumes using the NPM1 staining at these two stages. This quantification shows that nucleoli are bigger at HH12-13 than at HH18-19, and that at this latter stage, they are bigger in the ventral domain, which produces neurons, compared to the dorsal domain (**Fig.4D**). To further investigate the regulation of RiBi, we assessed the phosphorylation status of RPS6, a component of the 40S ribosomal subunit and a downstream target of mTORC1 and S6 Kinase signalling. (Liu and Sabatini, 2020). P-RPS6 is detected in a few cells in the ventral progenitor domain at HH12-13 (**Fig. 4B**), and in a higher proportion of ventral cells at HH18-19 (**Fig. 4C**), indicating that the mTORC1 pathway is also reinforced in the ventral neurogenic region, consistent with the increase in RiBi. Lastly, using transmission electron microscopy (TEM), we quantified the number of cytoplasmic ribosomes (most of them being translating ribosomes) in these same domains **(Fig4E)**. The average number of cytoplasmic ribosomes in progenitors was similar at HH12 and HH18-19, and did not differ significantly between the ventral and dorsal domains at HH18-19. (**Fig. 4E, 4F).** Hence, if ribosomes production is increased in nucleolus, it does not immediately lead to an increase of ribosomes in the cytoplasm. This result suggests that translation is not yet enhanced in the neurogenic ventral progenitors and that the increase in RiBi in the ventral region does not trigger an immediate change in the translation rate within this domain.

Altogether, these expression patterns reveal an unexpected dynamic pattern of RiBi during neurogenesis. At early stage, when progenitors mainly undergo proliferative divisions, RiBi levels are homogeneous and high. Then, the global RiBi level decreases upon neural tube maturation but is sustained in the neurogenic region compared to the proliferative one. The higher level of RiBi in progenitors of the neurogenic domain is associated with a higher level of the S6K triggered-mTORC1 pathway activity (**Fig.4G**). Moreover, the difference between the FBL and NPM1 patterns at HH18-19 reveals a specific level of regulation of later steps of ribosome assembly in neural progenitors as they engage toward differentiation.

### Sequential increases in RiBi, mTORC1 signalling and translation accompany neuronal commitment and differentiation

The temporal analysis of RiBi reveals that it is dynamically regulated during neurogenesis. Neural progenitors undergo two important switches during maturation, first committing to the neuronal fate and then differentiating into neurons (**Fig.5A**). To determine if the RiBi changes are linked to these particular switches, and if it consequently triggers enhanced translation rate, we deepened our analysis on the ventral domain of the cervico-brachial region at the HH18-19 stage. We used OLIG2 to label ventral neural progenitors, NEUROG2 to mark neuronal precursors committed to neuronal differentiation, and HUC/D or Islet1 to label neurons (**Fig. 5A**).

**Figure 5:**
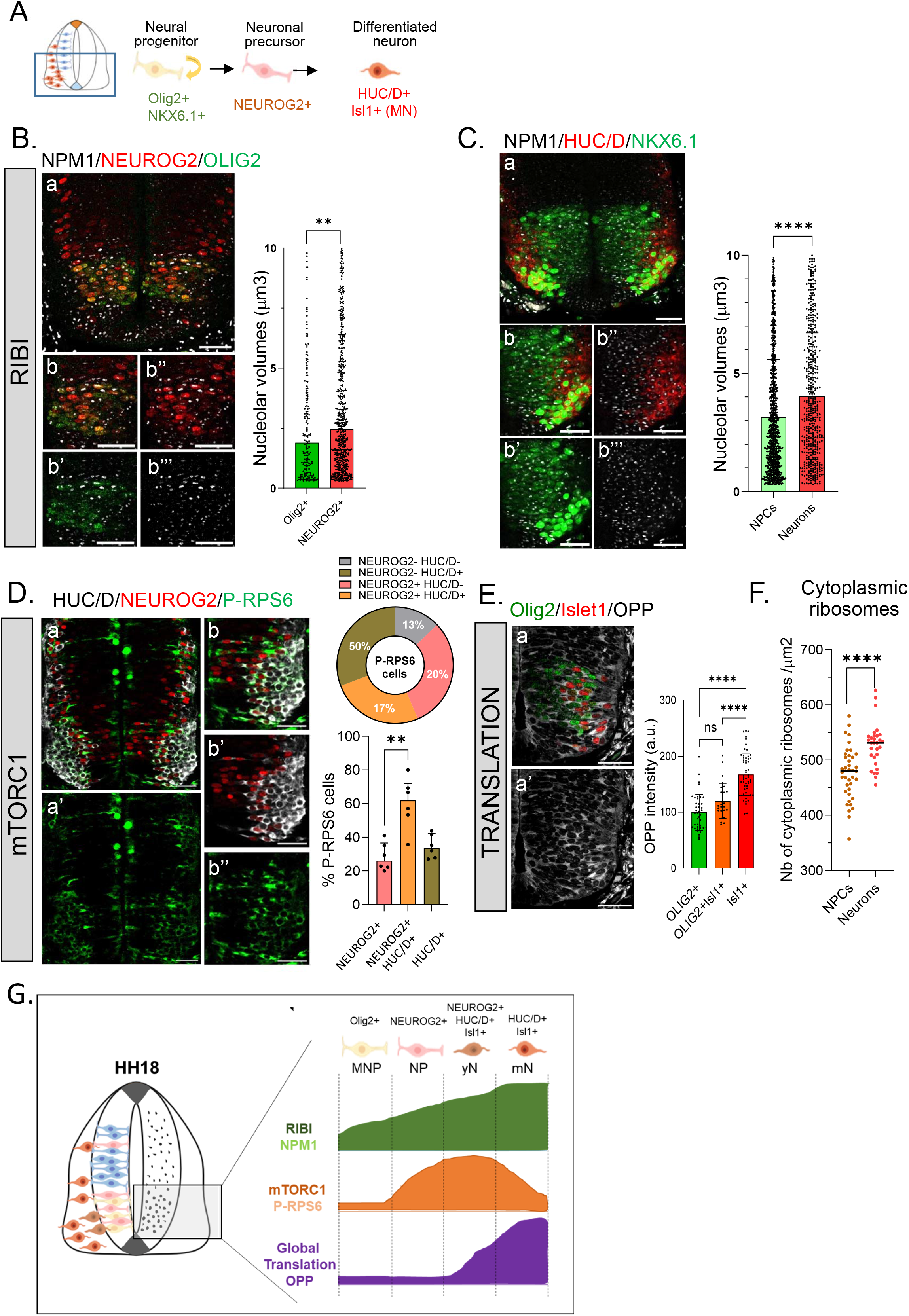
Sequential increases in RiBi, mTORC1 signaling and translation accompany neuronal commitment and differentiation. (A) Schematic representation of neural progenitor maturation stages. Ventral neural progenitors express Olig2 and/or NKX6.1, followed by a neuronal precursor state stained with NeuroG2 and Tis21. Differentiated neurons within the motor neuron domain are identified by HuC/D and Isl1 expression. (B) Left: Transverse sections of the HH18 chicken spinal cord stained for NPM1 (Grey), NEUROG2 (Red) and Olig2 (Green). a. large view of the ventral domain; b-b’’’ zoom up of half of the ventral region. Right: Nucleolar volumes (µm³) quantified from NPM1 immunostaining in (OLIG2+NEUROG2-) or (OLIG2+NEUROG2+) + (OLIG2-NEUROG2+) cells. Quantifications were performed across at least 3 embryos, with data expressed as median ± IQ range (each point representing an individual nucleolus). Mann-Whitney test, ** p< 0,01. Scale bars 30 µm. (C) Left: Transverse sections of the HH18 chicken spinal cord stained with NKX6.1 (green, ventral progenitors faint staining and V3 interneurons strong staining overlapping the HUC/D+ staining), HUC/D (red) and NPM1 (grey). Right: Quantification of nucleolar volumes from NPM1 immunostaining the ventral progenitors or neurons. Mann-Whitney test, **** p< 0,0001. Scale bars 30 µm. (D) Left: Transverse sections of the HH18 chicken spinal cord ventral domain, stained with HUC/D (Grey), NEUROG2 (Red) and P-RPS6 (Green). Right: Proportion P-RPS6 positive cells in (NEUROG2+; HUC/D-) or (NEUROG2+; HUC/D+) or (NEUROG2-; HUC/D+) cells. Quantifications were performed across at least 6 embryos, with data expressed as median ± IQ range (each point representing the percentage of p-RPS6 per cells on one section). Kruskal-wallis test, ** p< 0,01. Scale bars 30 µm. (E) Left: Close-up views of one side of the ventral domain of HH18 chicken spinal cord transverse sections showing O-propargyl-puromycin (OPP) incorporation after a 30 min labelling pulse prior to sample collection. Sections were subsequently stained for Olig2 (Green) and Islet1 (Red). Right: OPP intensity quantification in (OLIG2+; Islet1-), (OLIG2+; Islet1+) and (OLIG2-; Islet1+) cells. Scale bars 30 µm. Quantifications were performed across at least 3 embryos, with data expressed as median ± IQ range (each point representing the mean intensity of a unique cell). Kruskal-wallis test, **** p< 0,0001. (F) Quantitative analysis of cytoplasmic ribosomes abundance in progenitors and neurons in the ventral domain at HH18. Quantifications were performed across at least 3 embryos at each stage, with data expressed as median ± IQ range (each point representing the value of the counting in one cell). Mann-Whitney test, **** p< 0,0001. (G) Schematic representation of RiBi, mTORC1 activity and translation at different steps of neurogenesis in the ventral neural tube based on our data. MNP (motoneuron progenitor), NP (neuronal precursor), yN (young neuron) and mN (mature neuron). Ribi increases in MNP followed by a boost of mTORC1 activity in NP and an increased translation rate in neurons.

We first compared nucleoli volumes in OLIG2+ ventral progenitors and NEUROG2+ neuronal precursors. We observed that the nucleoli volume is increased in NEUROG2+ neuronal precursors (**Fig. 5B**). To confirm this result, we electroporated SOX2/TIS21 sensors. We observed that nucleolar volumes are significantly increased in neurogenic progenitors confirming an increase of RiBi when progenitors commit to differentiating divisions **(Fig. S5).** We then quantified nucleoli in NKX6.1 progenitors and HUC/D + neurons and observed a marked increase of RiBi in neurons (**Fig. 5C**).

We then analysed mTORC1 activity reflected by the P-RPS6 staining in the different neural populations. Among the P-RPS6+ cells of the ventral domain, only 13% are neural progenitors (NEURO2-HUC/D-). 20% are committed to neuronal differentiation (NEUROG2+HUC/D) and 67% are differentiating neurons (NEUROG2+HUC/D+ and NEUROG2-HUC/D+) (**Fig.5D**). Among the neurogenic populations, around 25% of neuronal precursors (NEUROG2+) are P-RPS6+ and this proportion clearly increases as these precursors engage toward differentiation (60% of NEUROG2+HUC/D+ cells). This proportion decreases again in differentiating neurons (25% in NEUROG2-HUC/D+ cells). Hence, mTORC1 activity is initiated in neural progenitors just prior neuronal commitment and maintained until they differentiate. We then sought to determine if this progressive increase of RiBi and mTORC1 signalling in committed neural precursors was correlated with higher translation rate in these cells. As a proxy of the translation rate, we quantified the amount of newly synthetized peptides following O-propargyl puromycin (OPP) incorporation *in ovo.* Revelation of the OPP staining in Olig2+ or Islet1+ cells clearly show that its intensity is stronger in Islet1+ expressing neurons, indicating a boost of translation in newly born neurons (**Fig. 5E**). Moreover, TEM quantification revealed a markedly higher density of cytoplasmic ribosomes in neurons than in progenitors, consistent with increased translational activity in the differentiating neurons **(Fig5F).**

Altogether, these analyses show that during the process of neurogenesis, RiBi increases in neural progenitors in differentiating domains. RiBi is then reinforced in neuronal precursors that activate the mTORC1 signalling as they engage toward differentiation (NEUROG2+HUC/D+ cells). This is followed by a boost of translational activity in newly born neurons (ISLET1+ cells) **(Fig5G).** These observations suggest that progenitors committed to differentiation progressively prepare the translational machinery to accommodate increased translation in neurons.

### Increasing mTOR activity is sufficient to trigger neuronal differentiation

The fact that RiBi and S6K activity increase in progenitors committed to neuronal differentiation opens the question of whether this increase is causal for neuronal differentiation. To assess this question we electroporated a phosphomimetic form of the S6K kinase (S6K-pMim) (Fishwick et al., 2010) (**Fig.6A**) or a CTL plasmid together with H2B-GFP at HH12-13. We first assessed S6K activity by analyzing p-RPS6 levels in electroporated cells 24 h after electroporation, and observed an increase in p-RPS6 in S6K p-MIM-electroporated cells, indicating enhanced S6K activity (**Fig.6B**). We then analyzed neuronal commitment and differentiation in the ventral and dorsal neural tube by quantifying the proportions of NEUROG2+, NEUROG2+HUC/D+ and HUC/D+ cells among electroporated cells. In the ventral neurogenic domain, increased S6K activity triggered a shift of NEUROG2+ cells toward the NEUROG2+HUC/D+ state, and a shift of NEUROG2+HUC/D+ cells toward the HUC/D+ state (**Fig.6C**). Each transition was also enhanced in the dorsal domain, where S6K-pMIM-electroporated cells showed a trend toward more NEUROG2+ cells together with higher proportions of NEUROG2+HUC/D+ and HUC/D+ cells. Notably, NEUROG2+ cells were decreased ventrally but increased dorsally, although these changes did not reach significance. Together with the changes in NEUROG2+HUC/D+ and HUC/D+ proportions, this opposite behaviour suggests that S6K promotes the neuronal precursor-to-neuron transition in the ventral domain, and the neural progenitor-to-neuronal precursor transition in the dorsal one. Altogether, these results indicate that increasing S6K activity in neural cells is sufficient to accelerate their progression to the next step of differentiation (**Fig.6D**).

**Figure 6:**
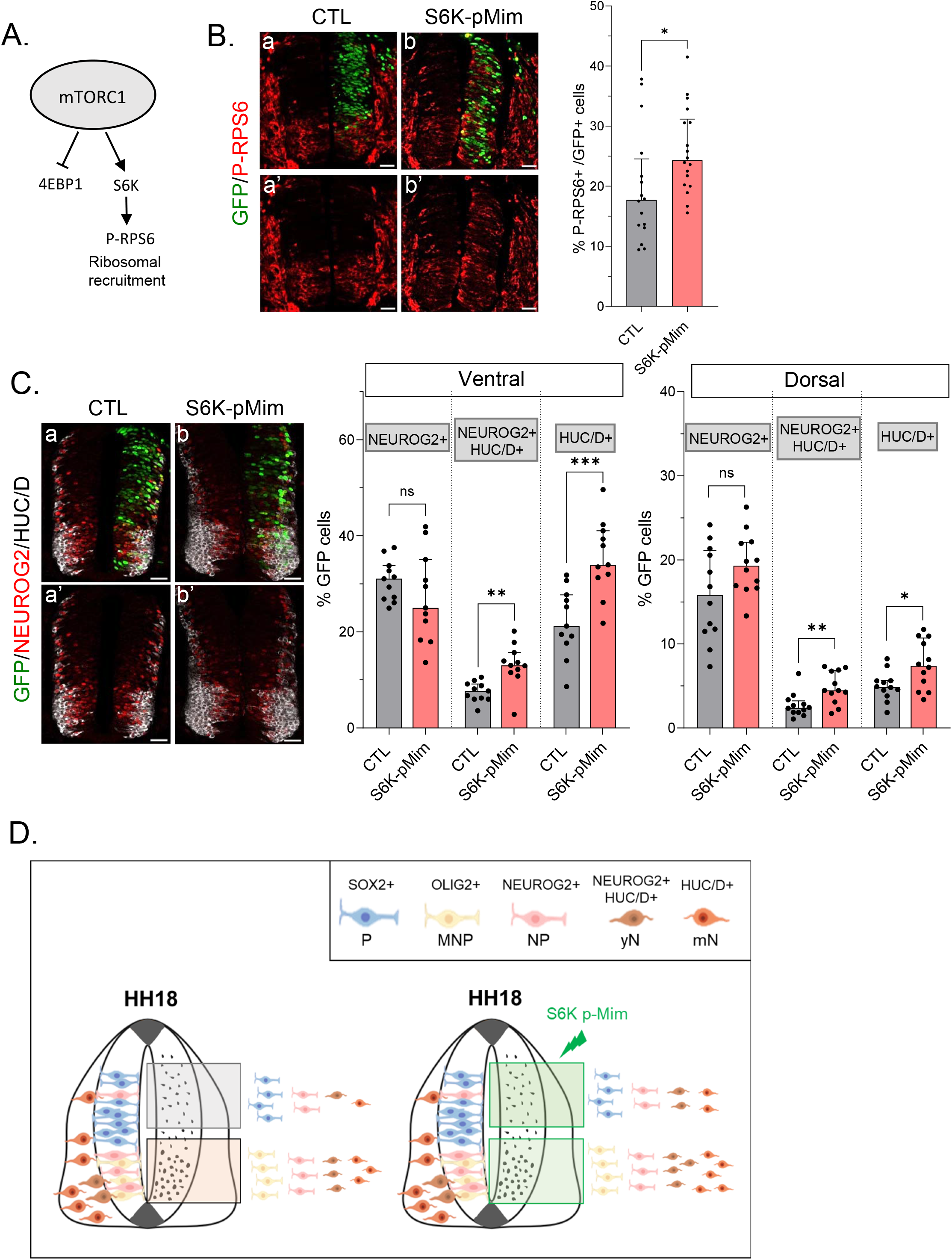
Increasing mTORC1 activity triggers a shift toward differentiation. (A) Scheme positioning S6 kinase and its target, P-RPS6, downstream of mTORC1 activity. (B) Left: Transverse sections of the HH18 chicken spinal cord electroporated with H2B-GFP (a,a’) or S6K-pMim (b,b’) and stained for P-RPS6 (red). Right: Quantification of the percentage of P-RPS6 cells among the GFP+ cells in the 2 conditions from 3 independent experiments. Each dot represents the mean value of the counting on 3 sections of one embryo. data expressed as median ± IQ range. Mann-Whitney test * p< 0,05. Scale bars 30 µm. (C) Left: Transverse sections of the HH18 chicken spinal cord electroporated with H2B-GFP alone (a,a’) or S6K-pMim (b,b’) and stained for NEUROG2 (red) and HUC/D (grey). Right: Quantification of the percentage of NEUROG2+, NEUROG2+HUC/D+ and HUC/D+ cells among the GFP+ cells in the 3 conditions from 3 independents experiments in the ventral or dorsal region. Each dot represents the mean value of the counting on 3 sections of one embryo. data expressed as median ± IQ range. Mann-Whitney test ***p< 0,001, **p< 0,01, *p< 0,05. Scale bars 30 µm. (D) Scheme recapitulating the effect of S6K-pMim on neuronal differentiation.

## DISCUSSION

In this study, using a RNA-Seq approach, we uncovered a link between the cell cycle machinery and ribosome biogenesis during spinal neurogenesis. We found that RiBi regulation is highly dynamic during spinal neurogenesis and that the cell cycle regulator CDC25B contributes to this regulation (**Fig.7**). We also show that increasing the mTORC1/S6K pathways in neural cells accelerate fate transition, from neural progenitors to neuronal precursors and neuronal precursors to neuron, thus leading to an increased neuronal differentiation rate. We propose a model in which, during neurogenesis, CDC25B coordinates cell cycle progression with RiBi in neurogenic progenitors, thereby increasing the rate of neuronal differentiation of their progeny (**Fig.7**).

**Figure 7:**
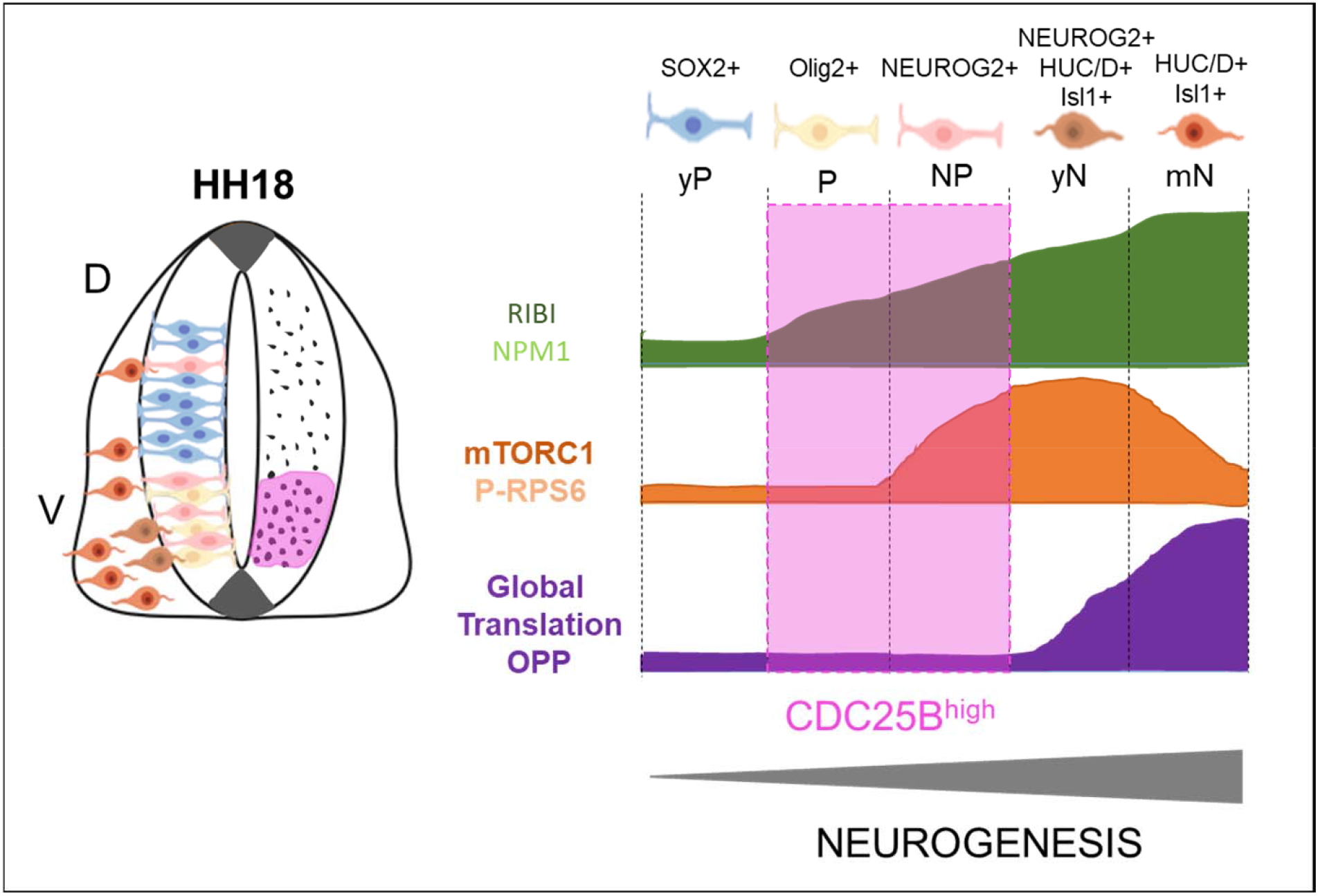
Model

Our RNA-seq data and functional analyses reveal that CDC25B regulates RiBi in spinal progenitors. We and others previously showed that CDC25B has two functions in neural progenitors: it modifies cell cycle kinetics, shortening G2 while indirectly lengthening G1, and it promotes neuronal differentiation (Peco et al., 2012; Bonnet et al., 2018; Roussat et al., 2023; Molina et al., 2022). Here, we show that increasing CDC25B activity upregulates genes involved in ribosome assembly and is associated with larger nucleoli, indicating that RiBi is stimulated downstream of this cell cycle regulator. In contrast, modulating the activity of CDK1, the main cell cycle target of CDC25B, had no detectable effect on RiBi. At first sight, this was unexpected, as CDK1 regulates RiBi and translation at multiple levels in other cellular models (Okuwaki et al., 2010; Haneke et al., 2020). However, in our experimental context, when shortening G2 with PD0166285, the only effect we observed was an increase in ventral neuronal differentiation without any effect in the dorsal neural tube. Yet, the extent of this modification was weaker than what we had observed with the CDC25B gain of function (Bonnet et al., 2018). This indicates that shortening G2 phase alone is not sufficient to drive efficient neuronal differentiation, and therefore the absence of RiBi changes upon PD0166285 treatment is not surprising. It further suggests that CDC25B regulates RiBi through its pro-neurogenic activity rather than through its effect on G2 length. We cannot fully exclude that the duration and or the dose of our treatment was not optimal and this could explain the mild effect on neurogenesis and the absence of effect on RiBi. Nevertheless, two non-exclusive mechanisms could account for the effect of CDC25B on RiBi. First, PD0166285 treatment did not alter G1 length, whereas CDC25B lengthens G1 phase (Molina et al., 2022). RiBi regulation by CDC25B could therefore be linked to G1 rather than G2 regulation, consistent with reports that ribosome production changes substantially during this phase (Iyer-Bierhoff and Grummt, 2019). Second, since part of the neurogenic activity of CDC25B is CDK-independent (Bonnet et al., 2018), RiBi regulation could involve the recruitment of other CDC25B partners. Further investigations will be necessary to clarify this point.

Our study also reveals a very dynamic regulation of RiBi and translation over the different steps of neurogenesis. At HH12-13, when progenitor domains are first specified along the dorso-ventral axis and, progenitors mainly perform proliferative divisions, nucleoli are large and uniformly distributed. At HH18-19, as motoneuron differentiation begins in the ventral neural tube, nucleoli display an overall reduction in size if compared to the HH12-13 stage, and a differential pattern of RiBi along the dorso-ventral axis, being markedly increased in ventral than in dorsal progenitors. This elevated RiBi is not accompanied by increased translation in progenitors themselves, as we did not detect higher OPP intensity or an increased cytoplasmic ribosome number in ventral progenitors compared to dorsal progenitors. Instead, we measure higher translation rate and ribosomes numbers in the newborn neurons derived from these progenitors. At both stages of our analysis, progenitors display equivalent proliferation rates, as do progenitors in the dorsal neural tube, where RiBi drops at the later stage (Kicheva et al., 2014; Molina and Pituello, 2017). The differences of RiBi activity that we observed between these domains are therefore unlikely to be driven by proliferation rate. These two stages correspond to the two-phases described by Kicheva and colleagues (Kicheva et al., 2014): an early phase of progenitor specification without differentiation, followed by a later phase in which progenitor identity remains stable while domain sizes are shaped by cell-type–specific differentiation rates. In light of these data, our finding may be interpreted as follows: during the early phase, high RiBi levels would support the translational demands required to produce proteins critical for neural progenitor specification. Later, RiBi would remain elevated specifically in regions engaging in differentiation to support a high differentiation rate supported by higher translation. Of note, an increase of RiBi does not necessarily result in a rise in translation as not all ribosomes are associated in polysomes (Kraushar et al., 2015). In addition, Beyond their structural role in ribosomes, some ribosomal proteins can contribute de the selective translation of specific mRNA subsets or perform additional extra-ribosomal functions (Shi et al., 2017; Simsek et al., 2017). Therefore, additional evidence will be needed to confirm that the increase in RiBi in spinal neurogenic progenitors directly triggers an increase of protein translation in newborn neurons.

Similar dynamics of RiBi and translation are described for stem cells differentiation. When NSCs and their progenitor cells are isolated at various stages of differentiation toward olfactory bulb interneurons, protein synthesis is found to rise and fall several times as neural differentiation progress (Baser et al., 2019; Saba et al., 2021). Corsini et al. also revealed that protein synthesis rates and ribosome biogenesis (RiBi) change dynamically as ESCs differentiate into embryoid bodies (Corsini et al., 2018). In the Drosophila ovary, germline stem cells (GSCs) display high RiBi levels that drop as their progeny begins to differentiate, concomitant with an increase in translation in these daughter cells (Sanchez et al., 2016b; Zhang et al., 2014; Breznak et al., 2023). A similar RiBi/translation shift profile is more largely documented in somatic and embryonic mammalian stem cells as they engage in differentiation (Blair et al., 2017; Ingolia et al., 2011; Sampath et al., 2008; Signer et al., 2014; Jarzebowski et al., 2018). Recent evidences also document that ribosome biogenesis and translation are not merely passive consequences of cell growth but are actively regulated during cortical neurogenesis and can influence cell fate decisions (Chau et al., 2018; Fujii et al., 2017; Harnett et al., 2022; Kondrashov et al., 2011). Thus, beyond its importance for stem cell maintenance and differentiation, the uncoupling between RiBi and translation levels maybe a general property of differentiating cells. To integrate these findings, Saba and colleagues propose a differentiation “switch” model: enhanced RiBi in progenitor cells would prime their progeny with an abundant ribosome pool, poised to support the burst of translation that accompanies differentiation. Our results in the developing spinal cord are fully consistent with this model.

Accompanying these dynamic RiBi and translation variations, the S6 kinase, downstream of the mTORC1 signaling, also shows an heterogenous activity as visualized with p-RPS6 expression pattern. Indeed, similar to RiBi, this activity emerges as progenitors become neurogenic, being restricted to a few ventral cells at HH12-13 and extending to a large proportion of neuronal precursors and young neurons in the ventral neurogenic domain by HH17-18. Since S6K regulates RiBi (Liu and Sabatini, 2020), it makes sense that its activity is reinforced concomitantly with higher RiBi levels. Additionally, we found that upregulation of mTORC1 signaling is sufficient to accelerate cell fate transitions over the different steps of neurogenesis, as shown by the rise in NEUROG2+HUC/D+ and HUC/D+ cells following S6K p-Mim electroporation in ventral domain and of NEUROG2+ and NEUROG2+HUC/D+ cells in dorsal ones. In coherence with this result, Fishwick and colleagues, shows that oppositely, neuronal production is reduced in the chick neural tube following exposure to rapamycin, an mTORC1 inhibitor, and that this same treatment repressed NEUROG2 expression (Fishwick et al., 2010). Together, these two sets of data indicate that mTORC1 signaling is both necessary and sufficient to trigger the differentiation of spinal neurons. Likewise, such positive control of mTORC1 signaling in driving progeny differentiation has been described in GSCs, ESCs and NSCs and several other stem cell types (Saba et al., 2021; Blair et al., 2017; Easley et al., 2010; Baser et al., 2019). Interestingly, Baser and colleagues describe that, if mTORC1 signaling is high in NSCs activated state, it transiently drops upon neuronal differentiation and cell cycle exit, triggering selective translational repression of stem cell identity factors (SOX2, PAX6) and components of the translation machinery, thereby driving the onset of differentiation (Baser et al., 2019). These finding further emphasize the very tight regulation of mTORC1 to orchestrate specific protein synthesis during the differentiation switch in different systems.

Interestingly, FBL and NPM1 show markedly different expression dynamics in spinal progenitors over time. FBL localizes to the fibrillar center and dense fibrillar component, driving early rRNA maturation, whereas NPM1 localizes to the granular component, where it governs later RiBi steps—40S/60S subunit assembly and nuclear export (Taha and Ahmadian, 2024; Zhang et al., 2024). Only NPM1, not FBL, displayed a dynamic expression pattern, suggesting that RiBi regulation occurs primarily at these final steps in our system. Indeed, it has been suggested that in multicellular organisms different steps of RiBi could be differentially regulated (Ni and Buszczak, 2023). Maintaining a homogenous level of FBL would allow to maintain basal level of ribosomal components that could be maturated, assembled and exported on demand to quickly increase protein production during the differentiation switch.

Finally, our WGCNA analysis of the RNA-Seq data uncovered a possible link between CDC25B and oxidative phosphorylation. Oxidative phosphorylation is the process by which cells produce most of their energy, in the form of ATP, using oxygen; it takes place in the inner mitochondrial membrane. This is particularly interesting, as we previously showed that mitochondria display differential morphology in neural progenitors versus neurons: they are short and thick in progenitors, supporting high glycolytic activity, and become filamentous in neurons, a morphological trait associated with oxidative phosphorylation (Mils et al., 2015). This switch toward oxidative phosphorylation in neurons has been shown, in other systems, to support the high energy demands of differentiating cells (Iwata et al., 2023). Given that mTORC1 signaling also controls mitochondrial oxidative function, this increase downstream of CDC25B could be directly linked to the rise of mTORC1 activity. This will be an interesting issue to address in future investigations.

## Material and methods

### Chicken embryos

Fertile chicken eggs at 38°C in a humidified incubator yielded appropriately staged embryos (Hamburger and Hamilton, 1992). Animal related procedures were performed according to EC guidelines (86/609/CEE), French Decree no. 97/748 and CNRS recommendation. Embryos were fixed 1h in paraformaldehyde prior sectioning on a vibratome for subsequent immunostaining.

### Electroporation and constructs

*In ovo* electroporation experiments were performed HH12-13 chicken embryos as described previously (Peco et al., 2012; Bel-Vialar et al., 2007). Gain-of-function experiments were performed using a vector expressing the human CDC25B (hCDC25B isoform 3) or CDC25B^ΔCDK^ (hCDC25B3ΔCDK) under the control of a cis regulatory element of the mouse Cdc25B called pccRE that reproduces the cell cycle regulated transcription of CDC25B and retain its function (Korner et al., 2001) (Bonnet et al., 2018). A control vector was generated with the βGal gene downstream of the pccRE. These plasmids were electroporated at 1.5 μg/μl together with an H2B-GFP at 0.8 μg /μl. The Sox2p-GFP and Tis21p-RFP sensors are a gift from E. Marti and used at 1 μg/μl and 0.5 μg/μl respectively. S6K-pMim is a gift from K.Storey (Fishwick et al., 2010).

### RNA-Seq Strategy

We overexpressed hCDC25B, hCDC25B^ΔCDK^ or the control vector in the neural tubes of HH12-13 chicken embryos together with an H2B-GFP to label electroporated cells. We harvested electroporated embryos 24h later and pooled 8 to 10 neural tubes for each biological repliquat. GFP positive cells were sorted using a FACS using a FACS ARIA Fusion machine (BD Biosciences) at the TRI-Genotoul platform (https://www.ipbs.fr/fr/tri-cytometrie/). RNA extraction was performed as described (Lacomme et al., 2012). We collected a mean of 60000 cells producing an average of 300 ng of RNA per sample. RNAs were checked for quality using Bioanalyser and sent to BGI for sequencing (https://services.bgi.com/bgi-rna-sequencing).

### RNA-Seq sequencing and DeSeQ analysis

RNA-Seq samples were sequenced using DNBSEQ-G50 sequencer (paired-end, 100bp long reads, 40M reads/sample) at BGI (www.BGI.com). The quality of the raw data and alignments were performed using the Galaxy web platform (https://galaxyproject.org/). The quality of each raw sequencing file (fastq) was verified with FastQC (https://www.bioinformatics.babraham.ac.uk/projects/fastqc/). Fastq were aligned the reference chicken genome GRCg6a (galGal6) using STAR aligner (STAR version 2.7). Processed bam files were then ordered and indexed using samtools version 1.12. Count read was computed using HT-seq version 0.13.5 on the Ensembl annotation (Gallus_gallus.GRCg6a.104_chr.gtf). Then the raw count table was filtered for lowly expressed genes (only gene expressed with at least 3 reads per sample in average were kept). Hierarchical clustering and heatmaps of differentially expressed genes were generated in R using the heatmap.2 command in ‘gplots’ package. Differential analysis (normalization and generation of log2FC and adjusted p-value) was applied using Bioconductor DESeq2 (version 1). We used principal component analysis (PCA) to cluster samples based on their normalized expression levels. RNA-Seq data have been submitted on the GEO database under accession number GSE352655. Gene set enrichment analysis was performed using Genecodis (https://genecodis.genyo.es/gc4) and STRING (https://string-db.org/).

### WGCNA analysis

The raw gene expression counts dataset (n = 15,497) for 15 samples (5 control CTRL, 5 hCDC25B^ΔCDK^, 5 hCDC25B) obtained from RNA-seq was normalised using the trimmed mean of M-values (TMM) method implemented in the edgeR package (Robinson et al., 2010) on the Asterics online platform (Maigné et al., 2023). This normalization aims to correct sequencing depth bias and make counts comparable between individuals. The filter threshold was set to 0.5, after which a log₂(x+0.5)-based transformation was applied.

Correlation analyses were performed using the WGCNA 1.74 package (Langfelder and Horvath, 2008) for weighted correlation network analysis with R software version 4.5.1 (“R: A language and environment for statistical computing – ScienceOpen,” n.d.). The analysis was conducted in accordance with the recommendations of the WGCNA developers ( (https://edo98811.github.io/WGCNA_official_documentation/). A sample clustering step was performed to detect possible outliers using the hclust function with the “average” method. One sample was excluded from the subsequent steps. The network was constructed by keeping the top 5000 genes with the highest variance. For the network construction, the similarity between pairs of genes was measured based on the Pearson coefficient. The first parameter determined was the soft power threshold for the sensitivity and the specificity of the pairwise connections (adjacency). This analysis of the network topology suggested choosing a soft-thresholding power of 4 to obtain a free-scale network. Then the module detection was performed with the following parameters: a minimum number of genes per module of 30 and a merge Cut Height of 0.25 to allow two modules to be merged if their correlation was greater than 0.75.

Next, it was possible to identify the module that was more significantly associated with the sample traits. Here, quantification of endogenousCDC25B, hCDC25B, and LacZ (used as control). A module eigengene was calculated for each module to create a single synthetic expression profile, and a correlation analysis was performed between each module eigengene and each trait.

### Immunostaining

Immunostaining was performed as described in (Lacomme et al., 2012). Vibratome sections were permeabilized in 0.5% triton X100 in PBS and blocked with 5% goat serum. For nucleolar staining, an additional permeabilization step with 2% triton X100 in PBS for 30 min was done prior the blocking step. We used primary antibodies against: NEUROG2 (guinea pig 1/30000, a gift from B.Novitch (Skaggs et al., 2011); OLIG2 (rabbit #AB9610, 1/300, Merck millipore); Islet1 (mouse #39.4D5, 1/6, hybridoma bank); NKX6.1 (rat #F55A10, 1/10, hybridoma bank); HUC/D (mouse #16A11, 1:500, Thermo Fisher Scientific); Fibrillarin (mouse #38F3, 1/300, Invitrogen); NPM1 (mouse #No-63, 1:300, Progen); P-RPS6 (rabbit # D57.2.2E, 1/300, Cell signalling technology). Secondary antibodies were from goat and coupled with Alexa Fluor 488, Alexa Fluor 555 or Alexa Fluor 647. Sections were counterstained with dapi to label nuclei.

### Quantification of RIBI by measuring the nucleolar volumes with IMARIS

Nucleolar volume provides a proxy for ribosome biogenesis levels, larger nucleoli reflecting an increased ribosome biogenesis (Baker, 2013; Sanchez et al., 2016b). We stained nucleoli with NPM1 on neural tube transversal sections. We then acquired optimised images with a 40X objective spanning 10 μm thickness with a leica SP8 confocal microscope. Nucleolar volumes were measured using the “Surface” module of the Imaris software according to published methods (Baker, 2013; Sanchez et al., 2016b; Shannon et al., 2018). We excluded volume <0.3 and >10 μm3 corresponding to debris or abnormal conglomerate structures.

### Cell cycle phase analyses

Changes in G2-phase length were estimated using the percentage of labelled mitoses (PLM) paradigm (Quastler and Sherman, 1959), as described in (Peco et al., 2012). Embryos were exposed to EdU for 90 min before fixation to label cells in S phase, and mitotic cells were then identified by immunostaining for phospho-histone H3 (PH3). The proportion of double-labelled cells (PH3⁺ EdU⁺) among all PH3⁺ cells was quantified; as the rate at which this proportion increases reflects G2-phase duration, it provides an estimate of changes in G2 length. EdU was detected using the Click-iT EdU Alexa Fluor 647 Imaging Kit (Invitrogen) according to the manufacturer’s instructions.

S-phase length (Ts) and total cell cycle length (Tc) were determined from the relative proportions of cells that had incorporated one or both thymidine analogues, EdU and BrdU (Martynoga et al., 2005). BrdU (500 µM) was first applied in ovo, followed 90 min later by EdU (500 µM, Invitrogen), and embryos were fixed 30 min after EdU administration. EdU was detected first, and BrdU was then detected using the G3G4 antibody (Developmental Studies Hybridoma Bank). The proportion of cells in G1 phase was determined as the fraction of Fucci-G1-positive cells (Sakaue-Sawano et al., 2008; Molina et al., 2022) among proliferating progenitors identified with anti-PCNA (Santacruz, # sc-56, clone PC10) immunostaining.

### OPP incorporation and detection

OPP quantification was performed according to the method described by (Liu et al., 2012). 200 μM of OPP was deposited on the heart of the chicken embryo. One hour later, the embryos were collected and processed either for immunostaining. The OPP signal was revealed by a fluorescent reaction using the Click-iT plus OPP protein synthesis kit (In Vitrogen). We quantified the fluorescent OPP signal in the cytoplasm of cells labelled with OLIG2 or Islet1 on 40X images using the ImageJ software. For each sample, the OPP intensity of 3 sections from 3 embryos was acquired and averaged using the average OPP intensity measured across 3 square regions of the section.

### Transmission electron microscopy

Tissues were fixed with 2,5% glutaraldehyde and 2% paraformaldehyde in Cacodylate buffer (0.1 M, pH 7.2, EMS, Hatfield, PA) overnight at 4°C, post-fixed with 1% OsO4 in the same buffer and finally treated for 1 h with 1% aqueous uranyl acetate. The samples were then dehydrated in a graded ethanol series and embedded in Epon (EMBed-812, EMS). After 48 h of polymerization at 60 °C, ultrathin sections (80 nm thick) were mounted on Formvar-coated copper slot grids. Finally, sections were stained with Uranyless and lead citrate (em-grade.com). Grids were examined with a TEM (Jeol JEM-1400, JEOL Inc, Peabody, MA, USA) at 80 kV. Images were acquired using a digital camera (Gatan Orius, Gatan Inc, Pleasanton, CA, USA) at a 15000X magnification (square images of around 1,8 μm2). Counting of the cytoplasmic ribosomes was performed using ImageJ. Briefly, images were binarized based on an intensity threshold and the segmented ribosomes (particules between 25-1000 nm2) were counted. Results are expressed in number of ribosomes/ μm2.

### Statistical analysis

For each experiment, we analysed at least 3 sections of three different embryos and experiments were repeated at least three times. Statistical analyses were performed using Graph Pad Prism 10 software. Appropriate statistical tests were selected based on the distribution of data, homogeneity of variances, and sample size. Statistical tests used and sample sizes are indicated in the figure legends. P-value are indicated as follow: *p_0.05, **p_0.01, ***p_0.001, ****p_0.0001.

## ACKNOWLEDGMENTS

We acknowledge the METi imaging facility, member of the national infrastructure France-BioImaging supported by the French National Research Agency (ANR-10-INBS-04) and the LITC imaging facility, member of Genotoul-TRI and the National Infrastructure France-BioImaging supported by the French National Research Agency (ANR-24-INBS-0005 FBI BIOGEN). We thank Marion Aguirrebengoa from the big-A facility at the CBI for her help processing and analysing sequencing data. We thank K.Storey, B.Novitch for the gift of antibody or DNA construct. We thank A.Davy, B.Benazeraf, E.Agius and V.Lobjois for constructive discussion and reading of the manuscript. Author contributions: S.O. and S.B.-V. designed research; S.O, L.L, S.B.V, S.L and C.P.C performed research; S.O. and S.B.-V. analyzed data; S.B.-V. wrote the paper, F.P. provided input to successive drafts of the manuscript and FP and S.B.-V. edited the paper. We acknowledge the Developmental Studies Hybridoma Bank, created by the NICHD of the NIH and maintained at The University of Iowa, Department of Biology, Iowa City, IA 52242, for supplying monoclonal antibodies. This work was supported by the Centre National de la Recherche Scientifique, Université de Toulouse, Ministère de L’Enseignement Supérieur et de la Recherche (MESR) et Agence Nationale de la Recherche (ANR-19-CE16-0006-01).

## Notes

https://www.ncbi.nlm.nih.gov/geo/query/acc.cgi?acc=GSE352655

### Competing Interest Statement

The authors have declared no competing interest.

