## Supplementary material for "CDC25B couples ribosome biogenesis to cell cycle kinetics to promote spinal cord neurogenesis": Figure S1

### Slide 1
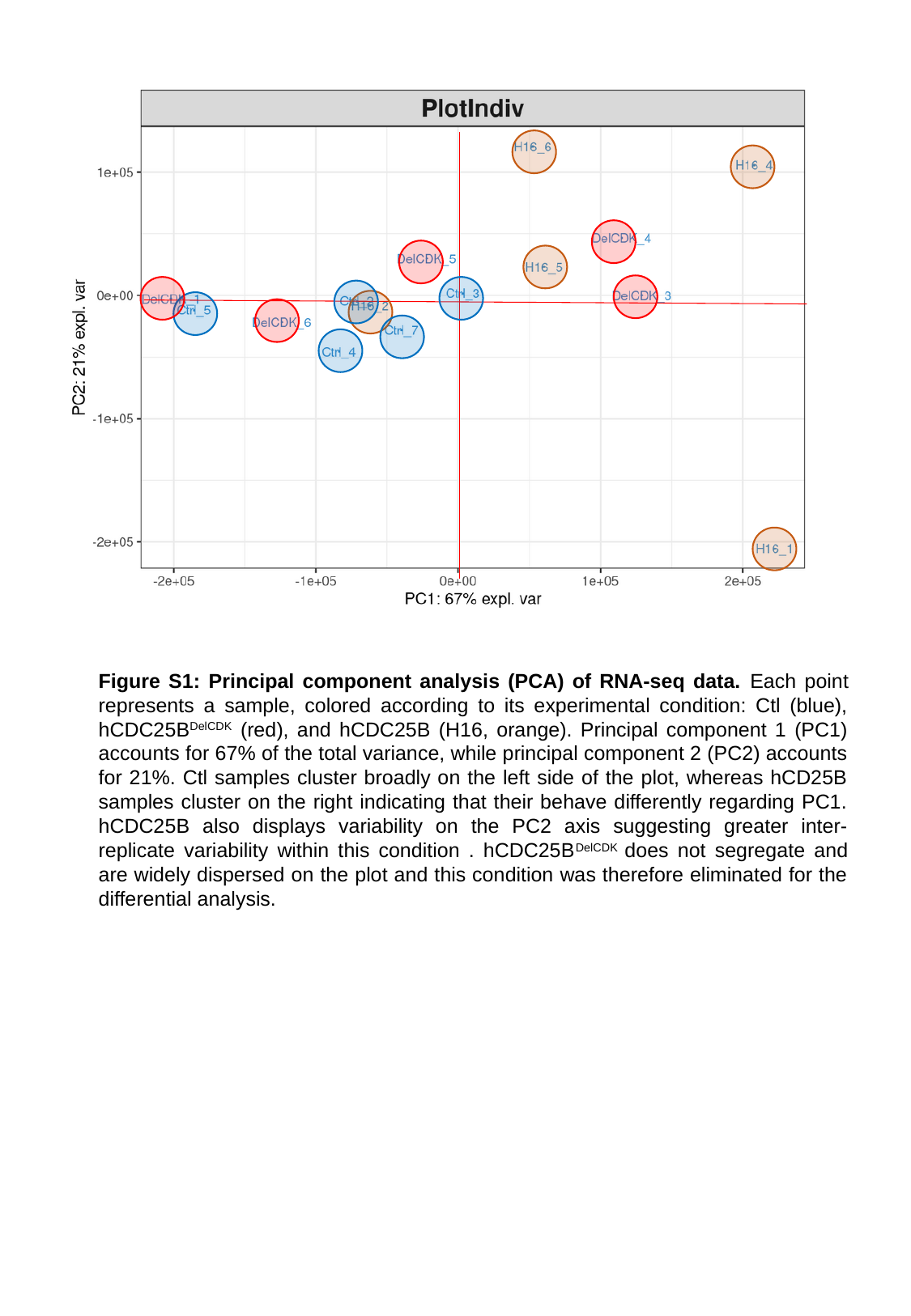

Figure S1: Principal component analysis (PCA) of RNA-seq data. Each point represents a sample, colored according to its experimental condition: Ctl (blue), hCDC25B^DelCDK^ (red), and hCDC25B (H16, orange). Principal component 1 (PC1) accounts for 67% of the total variance, while principal component 2 (PC2) accounts for 21%. Ctl samples cluster broadly on the left side of the plot, whereas hCD25B samples cluster on the right indicating that their behave differently regarding PC1. hCDC25B also displays variability on the PC2 axis suggesting greater inter-replicate variability within this condition . hCDC25B^DelCDK^ does not segregate and are widely dispersed on the plot and this condition was therefore eliminated for the differential analysis.
