## Supplementary material for "CDC25B couples ribosome biogenesis to cell cycle kinetics to promote spinal cord neurogenesis": Figure S2

### Slide 1
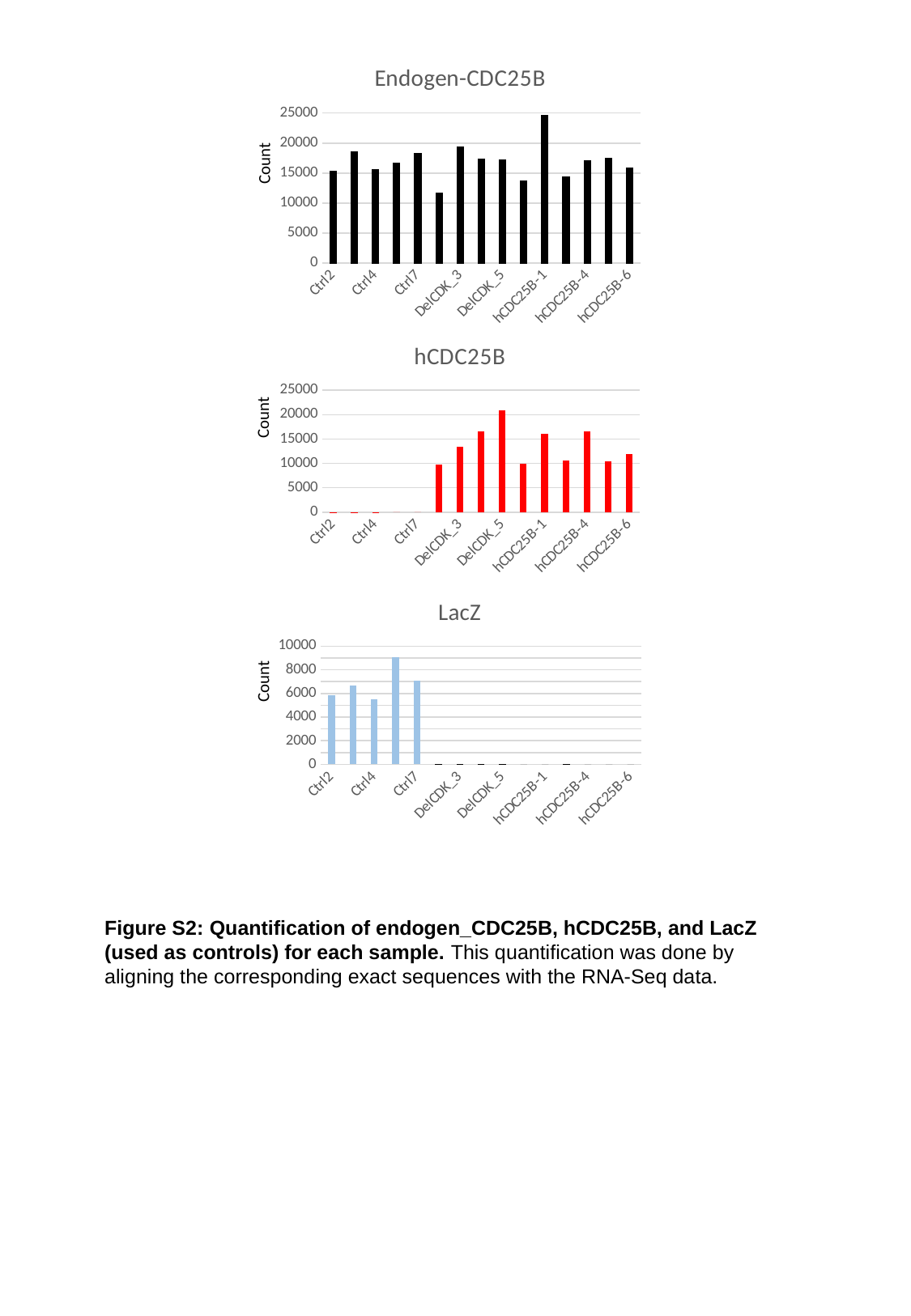

#### Chart: Endogen-CDC25B
| Category | Endogen_CDC25B |
|---|---|
| Ctrl2 | 15362.0 |
| Ctrl3 | 18492.0 |
| Ctrl4 | 15568.0 |
| Ctrl5 | 16662.0 |
| Ctrl7 | 18236.0 |
| DelCDK_1 | 11662.0 |
| DelCDK_3 | 19376.0 |
| DelCDK_4 | 17366.0 |
| DelCDK_5 | 17160.0 |
| DelCDK_6 | 13692.0 |
| hCDC25B-1 | 24600.0 |
| hCDC25B-2 | 14372.0 |
| hCDC25B-4 | 17100.0 |
| hCDC25B-5 | 17490.0 |
| hCDC25B-6 | 15818.0 |Count
#### Chart:
| Category | hCDC25B |
|---|---|
| Ctrl2 | 2.0 |
| Ctrl3 | 2.0 |
| Ctrl4 | 14.0 |
| Ctrl5 | 0.0 |
| Ctrl7 | 0.0 |
| DelCDK_1 | 9750.0 |
| DelCDK_3 | 13442.0 |
| DelCDK_4 | 16608.0 |
| DelCDK_5 | 20838.0 |
| DelCDK_6 | 9952.0 |
| hCDC25B-1 | 16036.0 |
| hCDC25B-2 | 10536.0 |
| hCDC25B-4 | 16608.0 |
| hCDC25B-5 | 10340.0 |
| hCDC25B-6 | 11952.0 |Count
#### Chart:
| Category | LacZ |
|---|---|
| Ctrl2 | 5876.0 |
| Ctrl3 | 6672.0 |
| Ctrl4 | 5522.0 |
| Ctrl5 | 9092.0 |
| Ctrl7 | 7058.0 |
| DelCDK_1 | 2.0 |
| DelCDK_3 | 6.0 |
| DelCDK_4 | 12.0 |
| DelCDK_5 | 2.0 |
| DelCDK_6 | 0.0 |
| hCDC25B-1 | 0.0 |
| hCDC25B-2 | 2.0 |
| hCDC25B-4 | 0.0 |
| hCDC25B-5 | 0.0 |
| hCDC25B-6 | 0.0 |Count
Figure S2: Quantification of endogen_CDC25B, hCDC25B, and LacZ (used as controls) for each sample. This quantification was done by aligning the corresponding exact sequences with the RNA-Seq data.
