## Supplementary material for "CDC25B couples ribosome biogenesis to cell cycle kinetics to promote spinal cord neurogenesis": Figure S3

### Slide 1
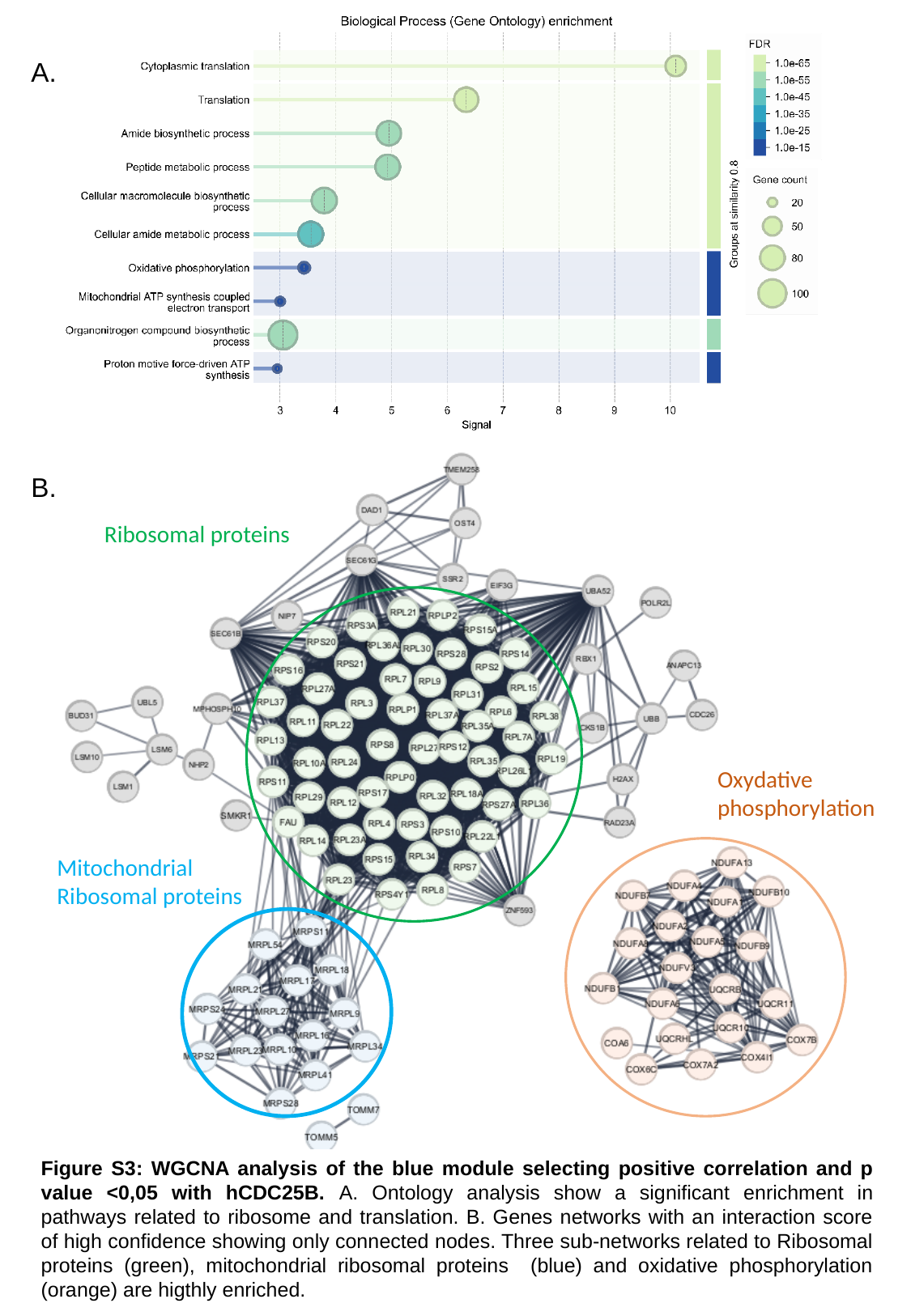

A.
B.
Ribosomal proteins
Oxydative phosphorylation
Mitochondrial Ribosomal proteins
Figure S3: WGCNA analysis of the blue module selecting positive correlation and p value <0,05 with hCDC25B. A. Ontology analysis show a significant enrichment in pathways related to ribosome and translation. B. Genes networks with an interaction score of high confidence showing only connected nodes. Three sub-networks related to Ribosomal proteins (green), mitochondrial ribosomal proteins (blue) and oxidative phosphorylation (orange) are higthly enriched.
Coll. Laurence Liaubet
