## Supplementary material for "CDC25B couples ribosome biogenesis to cell cycle kinetics to promote spinal cord neurogenesis": Figure S4

### Slide 1
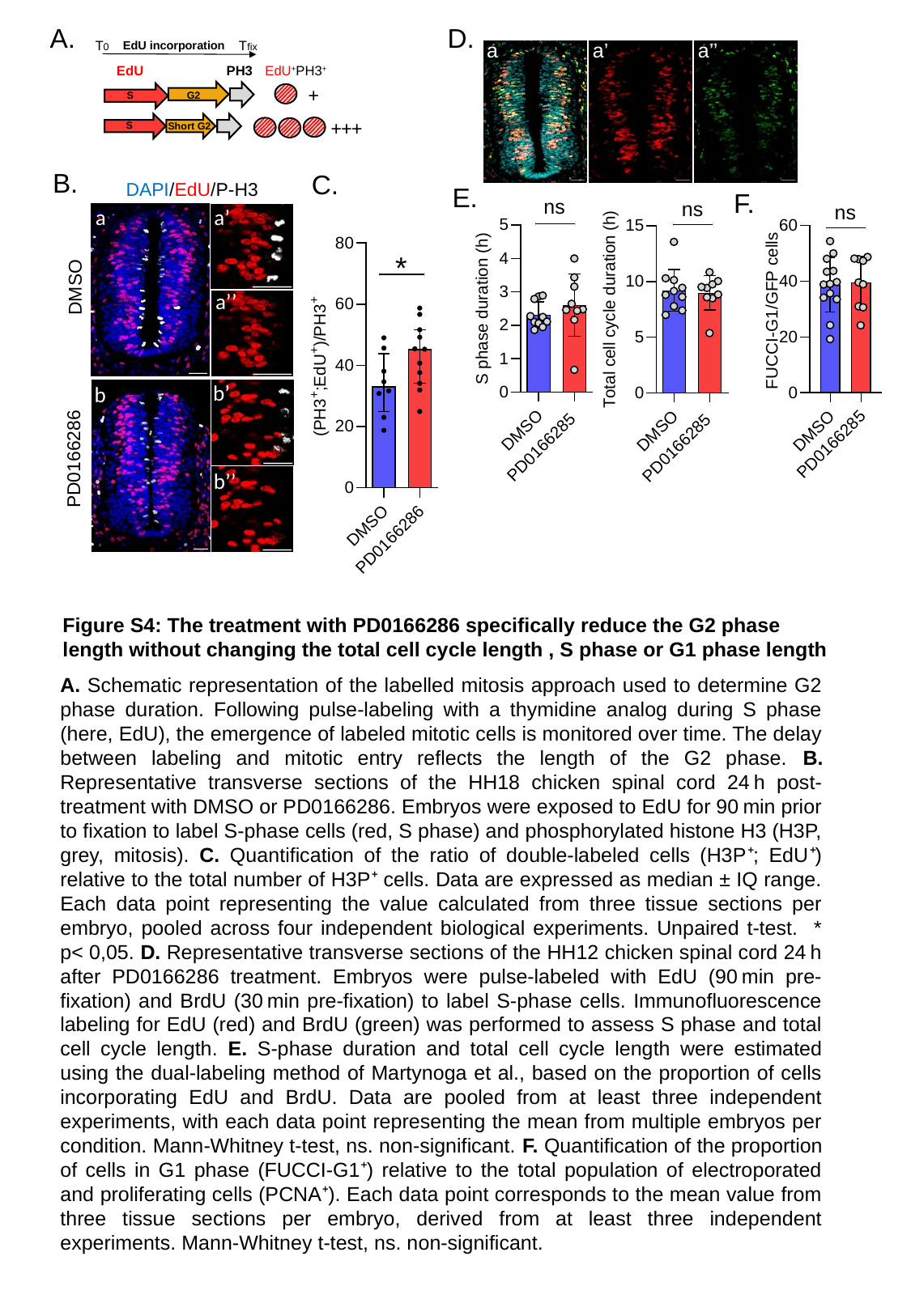

A.
D.
T0
Tfix
EdU incorporation
EdU
EdU^+^PH3^+^
PH3
+
G2
S
+++
S
Short G2
a
a’
a’’
B.
C.
DAPI/EdU/P-H3
E.
F.
a’
a
DMSO
a’’
b’
b
PD0166286
b’’
Figure S4: The treatment with PD0166286 specifically reduce the G2 phase length without changing the total cell cycle length , S phase or G1 phase length
A. Schematic representation of the labelled mitosis approach used to determine G2 phase duration. Following pulse-labeling with a thymidine analog during S phase (here, EdU), the emergence of labeled mitotic cells is monitored over time. The delay between labeling and mitotic entry reflects the length of the G2 phase. B. Representative transverse sections of the HH18 chicken spinal cord 24 h post-treatment with DMSO or PD0166286. Embryos were exposed to EdU for 90 min prior to fixation to label S-phase cells (red, S phase) and phosphorylated histone H3 (H3P, grey, mitosis). C. Quantification of the ratio of double-labeled cells (H3P⁺; EdU⁺) relative to the total number of H3P⁺ cells. Data are expressed as median ± IQ range. Each data point representing the value calculated from three tissue sections per embryo, pooled across four independent biological experiments. Unpaired t-test. * p< 0,05. D. Representative transverse sections of the HH12 chicken spinal cord 24 h after PD0166286 treatment. Embryos were pulse-labeled with EdU (90 min pre-fixation) and BrdU (30 min pre-fixation) to label S-phase cells. Immunofluorescence labeling for EdU (red) and BrdU (green) was performed to assess S phase and total cell cycle length. E. S-phase duration and total cell cycle length were estimated using the dual-labeling method of Martynoga et al., based on the proportion of cells incorporating EdU and BrdU. Data are pooled from at least three independent experiments, with each data point representing the mean from multiple embryos per condition. Mann-Whitney t-test, ns. non-significant. F. Quantification of the proportion of cells in G1 phase (FUCCI-G1⁺) relative to the total population of electroporated and proliferating cells (PCNA⁺). Each data point corresponds to the mean value from three tissue sections per embryo, derived from at least three independent experiments. Mann-Whitney t-test, ns. non-significant.
