## Supplementary material for "CDC25B couples ribosome biogenesis to cell cycle kinetics to promote spinal cord neurogenesis": Figure S5

### Slide 1
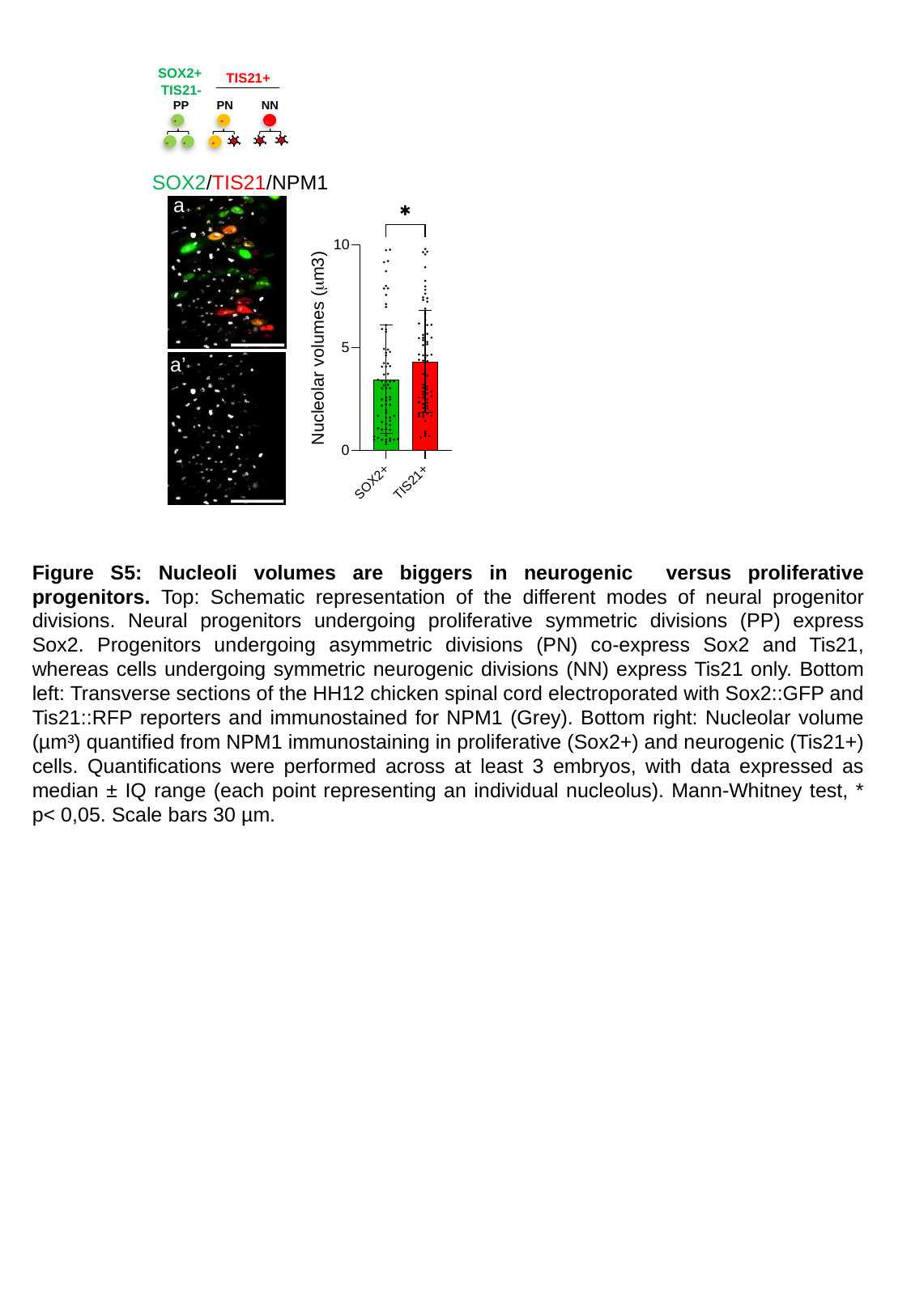

SOX2+
TIS21+
TIS21-
PP
PN
NN
SOX2/TIS21/NPM1
a
a’
Figure S5: Nucleoli volumes are biggers in neurogenic versus proliferative progenitors. Top: Schematic representation of the different modes of neural progenitor divisions. Neural progenitors undergoing proliferative symmetric divisions (PP) express Sox2. Progenitors undergoing asymmetric divisions (PN) co-express Sox2 and Tis21, whereas cells undergoing symmetric neurogenic divisions (NN) express Tis21 only. Bottom left: Transverse sections of the HH12 chicken spinal cord electroporated with Sox2::GFP and Tis21::RFP reporters and immunostained for NPM1 (Grey). Bottom right: Nucleolar volume (µm³) quantified from NPM1 immunostaining in proliferative (Sox2+) and neurogenic (Tis21+) cells. Quantifications were performed across at least 3 embryos, with data expressed as median ± IQ range (each point representing an individual nucleolus). Mann-Whitney test, * p< 0,05. Scale bars 30 µm.
